# Rare variant in intracellular loop-2 of the ghrelin receptor reveals novel mechanisms of GPCR biased signaling and trafficking

**DOI:** 10.1101/2025.03.11.642421

**Authors:** Elsa M. Balfe, Alexandre Torbey, Lara Kohlenbach, Jade A. Sency, Asuka Inoue, Lawrence S. Barak, Joshua D. Gross

## Abstract

G protein-coupled receptors (GPCRs) are the largest class of integral membrane proteins and the most common pharmaceutical drug target. Through allosteric coupling, GPCRs transduce extracellular stimuli into physiologically-relevant intracellular signaling cascades via G proteins and β-arrestin (βarr). The growth hormone secretagogue receptor (GHSR) is a rhodopsin-like, peptide hormone GPCR considered a promising target for both metabolic and neurological diseases. Here, by characterizing an ultra-rare coding variant in intracellular loop-2 (ICL2) of the GHSR, GHSR^L149P^ (L149P), we establish a unique role of ICL2 in GPCR biased signaling, lipid modulation, and intracellular trafficking. Using an array of bioluminescence resonance energy transfer (BRET)-based assays, we show that the natural L149P mutant exhibits (i) a constitutive plasma membrane [PM] expression bias, (ii) preferential partitioning away from cholesterol-enriched PM microdomains, (iii) enhanced agonist-directed endocytosis, and (iv) dramatic signaling bias towards βarr1/2 over Gα_q_, Gα_i/o_, and Gα_12/13_. Using a combination of pharmacological and genetic tools, we demonstrate that βarr1/2 recruitment to L149P requires G protein-coupled receptor kinase-2/3 (GRK2/3)-mediated phosphorylation, but it does not utilize protein kinase C (PKC), Gβγ-dependent GRK2/3 translocation, or Gα_i/o_, supporting a G protein-independent mechanism. Lastly, we found that βarr1/2 recruitment to L149P requires both GRK2/3 and GRK5/6, while the wild-type GHSR relies exclusively on GRK2/3, consistent with increased GRK6 pre-coupling to the L149P mutant. Collectively, our findings using a rare, natural variant reveal novel mechanisms of GPCR regulation that could be leveraged to improve personalized medicine and facilitate the rational design/discovery of GPCR^ICL2^-directed drugs.

**SIGNIFICANCE STATEMENT:** G protein-coupled receptors (GPCRs) are the largest class of membrane proteins and the most prevalent drug target class in medicine. However, the structural and mechanistic determinants of GPCR signaling and membrane compartmentalization remains incompletely understood. Here, we characterize an ultra-rare natural variant of the growth hormone secretagogue receptor (GHSR), L149P, that uncovers an unanticipated role for intracellular loop 2 (ICL2) in receptor biased signaling, intracellular trafficking, and membrane lipid modulation. The variant exhibits a strong bias for β-arrestin signaling independent of canonical G protein pathways, mediated instead by unique GPCR kinase interactions and a distinct subcellular distribution. These findings reveal previously unrecognized regulatory layers in GPCR structure/function and identify ICL2 as a promising target for designing precision pharmacotherapeutics.

## INTRODUCTION

G protein-coupled receptors (GPCRs) are the largest class of integral membrane proteins in the human genome and remain the most common pharmaceutical target ^1^. As allosteric protein complexes, GPCRs convert extracellular stimuli into intracellular signaling cascades via coupling to transducers; namely, G proteins and β-arrestin (βarr) ^2^. The evolutionary conservation and ubiquitous expression of GPCRs highlights how even rare, naturally-occurring GPCR mutations can precipitate clinically-relevant phenotypes ^3^. To enable adaptive biological responses, GPCRs adopt a multitude of conformational states that can asymmetrically engage G proteins or βarr, thereby leading to distinct physiological outcomes in a pathway-dependent manner (*i.e.*, biased signaling). To regulate these complex processes under dynamic environmental conditions, GPCRs harbor a diverse array of allosteric binding sites accessible to various ligands, regulatory proteins, and lipids. Accordingly, GPCR biased signaling can be engendered via ligand binding (biased ligands), receptor mutations (biased receptors), differential regulatory protein or lipid expression (systems bias), and/or differential receptor or transducer subcellular distribution (location bias) ^4^. Based on these principles, therapeutic interests have emerged around the promise of ‘biased’ pharmaceuticals that could enhance therapeutic efficacy while simultaneously reducing side effects, toxicity, or tachyphylaxis ^5^.

The growth hormone secretagogue receptor (GHSR) is the cognate target of the gut-derived, peptide hormones ghrelin (endogenous agonist) and liver-expressed antimicrobial protein-2 (LEAP-2; endogenous antagonist). The GHSR is a rhodopsin-like, Class A GPCR that preferentially couples to Gα_q_ and promiscuously engages members of the Gα_i/o_ and Gα_12/13_ subfamilies ^6^. Upon activation, the GHSR is phosphorylated by intracellular kinases, including Gα_q_-dependent protein kinase C (PKC) and Gβγ-dependent G protein-coupled receptor kinase 2/3 (GRK2/3), leading to βarr recruitment and subsequent desensitization, internalization/endocytosis, endosomal trafficking, and βarr-mediated signaling ^7^. The GHSR is expressed both centrally and peripherally, and it serves as an integral rheostat of energy balance, reward, neuronal health, and stress/affective behavior ^8^. Accordingly, the GHSR is a rational pharmaceutical drug target for several metabolic and neurological diseases ^9^. The GHSR can stabilize conformations conducive to biased signaling, suggesting that the discovery of biased GHSR ligands or receptors is mechanistically feasible ^6, 9–14^. While several GHSR X-ray crystallography and cryogenic electron microscopy (cryo-EM) structures have illuminated the molecular basis of ligand binding and receptor activation ^15–18^, the mechanisms and conformational signatures governing GHSR biased signaling and intracellular trafficking have not been elucidated fully ^9^.

Plasma membrane (PM) lipids are important allosteric regulators of GPCR expression/stability, ligand binding, and signaling ^19^. Ligand binding and downstream G protein activation of the GHSR are dependent on cholesterol and sphingolipids ^20^—key constituents of PM lipid raft microdomains ^21^. Moreover, cholesterol has been resolved bound to transmembrane clefts in at least four-out-of-seven GHSR structures ^22^, highlighting its importance in GHSR structure/function. While these effects are well-established, little is known about how membrane lipids affect GHSR partitioning within different subcellular compartments, and how these processes influence transducer coupling, downstream signaling, or receptor trafficking. Given that the GHSR is highly sensitive to diet-derived lipids and metabolic states impacting PM lipid composition ^23^, the cellular lipid microenvironment could play an important role in GHSR-related physiology. Recent studies suggest that GPCRs can be subdivided by their preference for specific GRK subtypes (GRK2/3-*versus* GRK5/6-preferring), and that these intrinsic features determine receptor biased signaling ^24–26^. For example, βarr-biased ligands (*e.g.*, TRV027 at angiotensin II type-1 receptor ^26^) and intrinsically βarr-biased receptors (*e.g.*, atypical chemokine receptors ^27^) preferentially utilize plasma membrane-anchored GRK5/6 for βarr recruitment, independently of Gβγ-mediated GRK2/3 translocation from the cytosol to the PM ^25^. While this is logical, as βarr-biased signaling should not produce free Gβγ (no G protein activation), a subset of GPCRs have been shown to recruit GRK2/3 and/or βarr in a Gβγ-independent manner ^28^. To our knowledge, a systematic analysis of GRK-dependent GHSR signaling has not been performed previously.

In this study, we identify and characterize an ultra-rare, coding variant in intracellular loop-2 (ICL2) of the GHSR, Leu149Pro (L149P). The Leu149^ICL2^ site (Ballesteros-Weinstein number 34.51) is +7 amino acids downstream of the highly conserved E/DRY motif, and we have shown previously that non-natural GHSR^Leu149^ mutations regulate receptor bias and trafficking ^10, 14^. Furthermore, Leu149 is situated within the short GHSR-ICL2 α-helix, which is critical for G protein activation ^29^. Structural data reveal that the short ICL2 α-helix is present in five-out-of-seven GHSR structures resolved to-date ^13–16^; thus, we hypothesized that mutations in Leu149^ICL2^ disrupt the ICL2 conformations/stability and thereby, exert pronounced effects on receptor expression/function.

Here, we found that the natural L149P mutant exhibits a dramatic expression and signaling bias, as well as altered membrane lipid modulation and intracellular trafficking; including, (i) increased constitutive PM expression relative to endosomes, (ii) reduced clustering within cholesterol-enriched, lipid raft-like PM microdomains, (iv) enhanced agonist-directed endosomal translocation despite reduced endocytosis from lipid rafts, (v) βarr1/2-biased signaling over multiple G protein subfamilies, (vi) G protein-independent βarr1/2 and GRK2 recruitment, and (vii) a GRK-subtype class switch from GRK2/3 dependence to GRK2/3/<u>5/6</u> dependence associated with increased GRK6 precoupling. Collectively, our results using an ultra-rare, natural GHSR variant support that the ICL2 conformational states govern GHSR biased signaling, lipid-dependent subcellular distribution, and intracellular trafficking.

## RESULTS

### The ultra-rare, natural variant — GHSR^L149P^ — exhibits altered subcellular distribution and trafficking in a cholesterol-dependent manner

Our prior work shows that contiguous residues in GHSR^ICL2^—located +6 and +7 amino acids downstream of the E/DRY motif (Pro148^34.50^ and Leu149^34.51^, respectively)—regulate receptor bias and trafficking ^10, 14^. To determine if natural mutations exist in these residues, we searched public genomic databases (NCBI Database for Single Nucleotide Polymorphisms [dbSNP]; build 156, released September 21^st^, 2022; https://www.ncbi.nlm.nih.gov/snp/rs1390872592) and identified Leu149>Pro (rs1390872592; **Figure 1a**), an ultra-rare nonsynonymous coding variant (reported allele frequency = 1/250,994). We did not identify natural variants in the upstream Pro148^34.50^ site, perhaps due to its high conservation across rhodopsin-like GPCRs ^30^.

**Figure 1.**
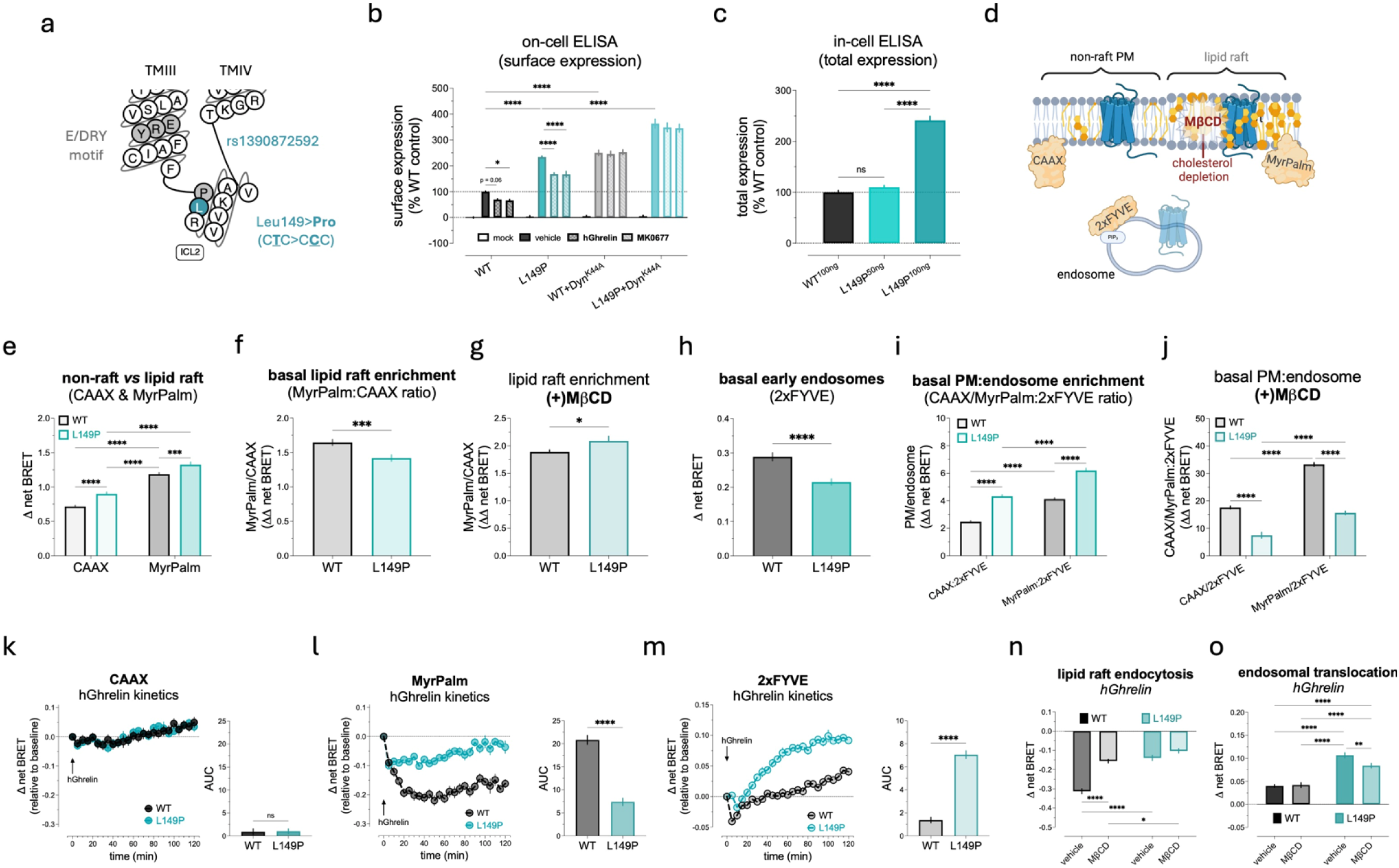
Subcellular distribution, PM partitioning, and trafficking of the natural L149P mutant. **(a)** GHSR Snake plot; E/DRY motif in TMIII (grey), Pro148^ICL2^ (grey), and Leu149^ICL2^ (teal). Non-synonymous SNP in Leu149 codon (C<u>T</u>C>C<u>C</u>C) resulting in the L149P missense mutation. **(b)** Basal and agonist (1μM)-stimulated (45-minutes, 37°C) surface expression (on-cell ELISA) in cells transiently expressing GHSR-WT^100ng^ (black/grey) or L149P^100ng^ (teal) ± Dyn^K44A^, analyzed by two-way ANOVA and Holm-Sidak’s post-hoc test. **(c)** Total receptor expression (in-cell ELISA) in cells transiently expressing WT^100ng^ (black), L149P^50ng^ (light teal), or L149P^100ng^ (dark teal), analyzed by one-way ANOVA. **(d)** bBRET sensors; mVenus-tagged CAAX (non-raft PM), MyrPalm (lipid raft PM), and 2xFYVE (early endosomes). MβCD (red) was used to deplete PM cholesterol. **(e)** Basal non-raft (CAAX) and lipid raft (MyrPalm) localization with WT^RLuc8^ (grey) or L149P^RLuc8^ (teal), analyzed by two-way ANOVA and Tukey’s post-hoc test. **(f)** Basal lipid raft enrichment calculated using the ratio of MyrPalm:CAAX (ΔΔ net BRET), derived from *Panel e*, then analyzed by *t*-test. **(g)** Basal lipid raft enrichment following MβCD pretreatment (3mM, 30-minutes, 37°C), analyzed by *t*-test. **(h)** Basal early endosomal (2xFYVE) localization, analyzed by *t*-test. **(i)** Basal PM enrichment over endosomes, derived from *Panels e* (CAAX and MyrPalm) and *Panel h* (2xFYVE), calculated using the ratio of CAAX:2xFYVE or MyrPalm:2xFYVE (ΔΔ net BRET), then analyzed by two-way ANOVA and Tukey’s post-hoc test. **(j)** Basal PM enrichment over endosomes following MβCD pretreatment, then analyzed by two-way ANOVA and Tukey’s post-hoc test. Kinetics (*left*) and area under the curve (AUC; *right*) of hGhrelin (1μM)-induced **(k)** CAAX, **(l)** MyrPalm, and **(m)** 2xFYVE localization with WT^RLuc8^ (black) or L149P^RLuc8^ (teal); AUC was evaluated statistically by *t*-test. hGhrelin (1μM)-induced **(n)** endocytosis from lipid rafts (MyrPalm) and **(o)** endosomal translocation (2xFYVE) over two-hours following 30-minutes pretreatment with vehicle or MβCD, assessed by two-way ANOVA and Tukey’s post-hoc test. All data are expressed as the mean ± SEM from ≥3 independent experiments in triplicate or quadruplicate.

We first tested the basal cell surface expression and agonist-induced internalization of the L149P mutant upon transient expression in HEK293/T cells using on-cell enzyme-linked immunosorbent assay (ELISA). Consistent with our previously characterized GHSR-Leu149^ICL2^ mutants ^10, 14^, the basal surface expression of L149P was increased by >2-fold relative to the wild-type GHSR (WT) (**Figure 1b**). Both the WT and L149P receptors underwent agonist-induced internalization following stimulation with human acyl-ghrelin (ghrelin) or the synthetic unbiased agonist MK0677 (**Figure 1b**); however, ghrelin-stimulated WT receptor internalization did not reach statistical significance (p=0.06). The vehicle-adjusted magnitude of ghrelin- and MK0677-induced L149P internalization was comparable to the WT receptor (28% versus 30-34%, respectively) (**Supplementary Figure 1**). To determine if increased L149P surface expression was due to altered constitutive, dynamin-dependent internalization (*i.e.*, ‘receptor cycling’), we co-expressed a dominant negative dynamin mutant (Dyn^K44A^) in the same experiments. Dynamin inhibition markedly increased the basal surface expression of both the WT and L149P receptors, albeit to a greater extent for the WT (250% relative to WT-pcDNA) compared to L149P (129% relative to L149P-pcDNA) (**Figure 1b**). Notably, basal WT-GHSR surface expression when co-expressed with Dyn^K44A^ was remarkably similar to L149P at baseline, supporting that reduced constitutive L149P internalization/cycling may account for its enhanced surface expression. The co-expression of Dyn^K44A^ completely blocked ghrelin and MK0677-induced internalization of the WT and L149P mutant, indicating that both receptors undergo dynamin-dependent, agonist-directed endocytosis.

To determine total receptor expression, we performed in-cell ELISA assays upon transient expression of L149P at an equal (1:1) or surface expression-adjusted (1:2) mass relative to the WT receptor. Expression of L149P at a 1:2 ratio produced similar total receptor expression; whereas, a 1:1 ratio of L149P-to-WT produced >2-fold L149P total expression relative to the WT receptor (**Figure 1c**), similar to its surface expression (see **Figure 1b**). Thus, in all subsequent experiments, we transfected an expression-adjusted ratio of L149P-to-WT receptor (1:2).

We next evaluated PM partitioning and endosomal trafficking of the natural L149P mutant using a bystander bioluminescence energy resonance transfer (bBRET)-based approach; including, the mVenus-tagged (i) non-lipid raft-targeting PM sensor CAAX [prenylation CAAX box of KRas], (ii) the lipid raft-targeting PM sensor myristoyl-palmitoylated (MyrPalm), and (iii) the early endosome-targeting sensor 2xFYVE [zinc finger domain of endofin], which binds phosphatidylinositol 3-phosphate [PIP_3_] (**Figure 1d**). Relative to WT, the L149P mutant had increased PM localization within both non-raft (CAAX) and lipid raft (MyrPalm) microdomains at baseline (**Figure 1e**). However, lipid raft enrichment (MyrPalm:CAAX ratio) of L149P was reduced relative to WT (**Figure 1f**), supporting that the L149P mutant preferentially partitions into non-raft PM microdomains. To confirm these effects, we pretreated cells with methyl-β-cyclodextrin (MβCD)—a membrane cholesterol chelator—then re-evaluated PM microdomain partitioning. MβCD pretreatment reversed the reduction in L149P lipid raft enrichment, such that L149P exhibited increased partitioning into MyrPalm-positive microdomains following cholesterol depletion (**Figure 1g**).

Additionally, basal L149P localization in early endosomes (2xFYVE) was markedly reduced relative to the WT receptor (**Figure 1h**). By comparing the ratio of non-raft PM (CAAX) and lipid raft PM (MyrPalm) to endosomal (2xFYVE) localization, we found that L149P displays a constitutive PM expression bias relative to endosomes (**Figure 1i**), consistent with enhanced basal surface expression and reduced receptor recycling measured by ELISA (see **Figure 1b-c**). Furthermore, MβCD pretreatment reversed the L149P mutant’s PM expression bias (**Figure 1j**), supporting that altered L149P receptor subcellular distribution is cholesterol-dependent.

To test agonist-induced endocytosis between each subcellular compartment, we stimulated bBRET sensor-expressing cells with a saturating concentration of ghrelin (1μM) and measured the change in CAAX, MyrPalm, and 2xFYVE localization with the WT or L149P receptors over two-hours. Both WT and L149P receptors displayed minimal endocytosis from non-raft (CAAX) microdomains and their magnitudes did not differ (**Figure 1k**). In contrast, agonist-induced WT receptor endocytosis from lipid rafts (MyrPalm) was robust, while lipid raft-mediated L149P endocytosis was significantly decreased relative to WT (**Figure 1l**). Conversely, both the rate and magnitude of ghrelin-induced L149P endosomal translocation was markedly increased relative to WT (**Figure 1m**), consistent more with a Class B-like trafficking profile. Upon MβCD pretreatment, ghrelin-induced WT receptor endocytosis from lipid rafts was reduced, while L149P endocytosis was unaffected (**Figure 1n**). Notably, WT receptor endocytosis from lipid rafts was reduced to a level similar to the L149P mutant under cholesterol-replete (vehicle-treated) conditions (**Figure 1n**), suggesting that the reduction in lipid raft-mediated L149P endocytosis (see **Figure 1l**) is related to membrane cholesterol levels. Furthermore, ghrelin-induced endosomal translocation of the L149P mutant, but not WT, was significantly reduced upon MβCD pretreatment (**Figure 1o**), suggesting that increased agonist-directed L149P endosomal translocation (see **Figure 1m**) is, at least partially, cholesterol-dependent.

Collectively, our results in **Figure 1** support that the L149P mutant exhibits a cholesterol-dependent (i) PM expression bias due to attenuated constitutive internalization/cycling, (ii) reduction in lipid raft clustering and preferential partitioning into non-raft PM microdomains, (iii) reduction in lipid raft-mediated, agonist-directed endocytosis, and (iv) enhancement in agonist-directed endosomal translocation. For all descriptive statistics and statistical analyses in **Figure 1**, see **Supplementary Table 1**.

### Basal and agonist-stimulated βarr1/2 recruitment to the L149P mutant is dynamin- and cholesterol-dependent

Non-natural mutations in GHSR-Leu149^ICL2^ produce βarr-bias ^10, 14^; thus, we next evaluated human βarr1/2 (hβarr1/2) recruitment to L149P using a BRET-based approach. Basal βarr1 recruitment (**Figure 2a**) to L149P was similar to the WT receptor (**Figure 2b**). Concentration-response (C/R) analysis of ghrelin-induced βarr1 recruitment revealed that ghrelin potency (EC_50_) at L149P was comparable to WT, while maximal efficacy (E_max_) was reduced by 53% (**Figure 2c**). Kinetic analyses showed that a saturating concentration of ghrelin (1μM) stimulated βarr1 recruitment to L149P at a rate similar to the WT receptor (**Figure 2d**). Similar to ghrelin, MK0677 evoked βarr1 recruitment to L149P with equal potency and reduced efficacy (57%) relative to WT (**Figure 2e**), as well as similar βarr1 association kinetics (**Figure 2f**). Statistical evaluation of ghrelin and MK0677 E_max_ showed that βarr1 recruitment efficacy at L149P was reduced to similar extents for both agonists relative to their respective WT controls (**Figure 2g**).

**Figure 2.**
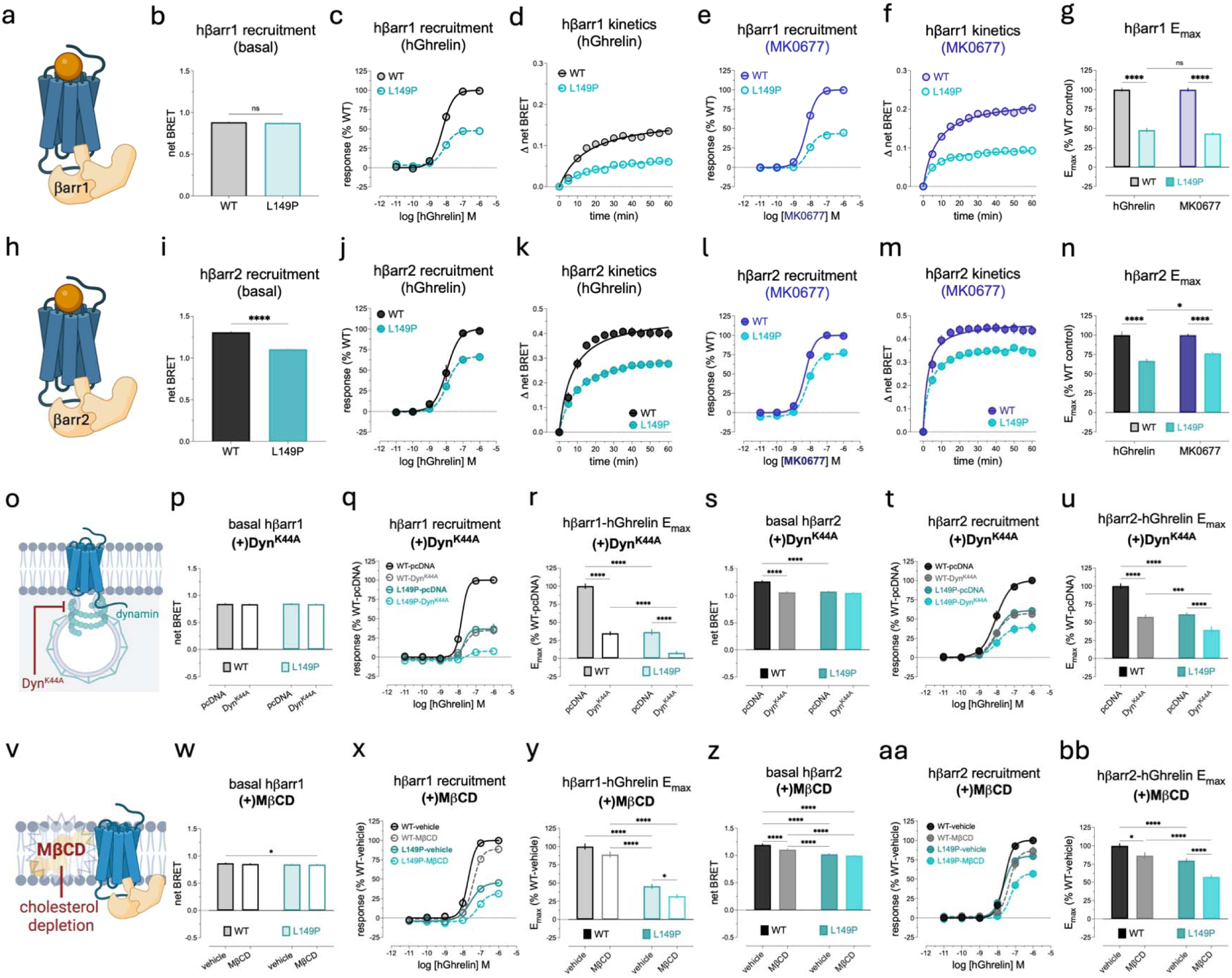
Basal and agonist-stimulated βarr1/2 recruitment to L149P is partially dynamin- and cholesterol-dependent. **(a)** βarr1 (light orange) coupling to GHSR (blue). **(b)** Basal hβarr1 recruitment to WT (grey) and L149P (light teal) expressed as absolute net BRET. **(c)** hGhrelin (C/R)-induced hβarr1 recruitment. **(d)** hGhrelin (1μM)-induced hβarr1 recruitment kinetics, derived from *Panel c*. **(e)** MK0677 (C/R)-induced hβarr1 recruitment to WT (light blue) and L149P (light teal). **(f)** MK0677 (1μM)-induced hβarr1 recruitment kinetics, derived from *Panel e*. **(g)** hGhrelin (*left*) and MK0677 (*right*) E_max_ for hβarr1 recruitment, derived from data in *Panel c* and *Panel e*, respectively. **(h)** βarr2 (light orange) coupling to GHSR (blue). **(i)** Basal hβarr2 recruitment to WT (black) and L149P (dark teal). **(j)** hGhrelin (C/R)-induced hβarr2 recruitment. **(k)** hGhrelin (1μM)-induced hβarr2 recruitment kinetics, derived from *Panel j*. **(l)** MK0677 (C/R)-induced hβarr2 recruitment to WT (dark blue) and L149P (dark teal). **(m)** MK0677 (1μM)-induced hβarr2 recruitment kinetics, derived from *Panel l*. **(n)** hGhrelin (*left*) and MK0677 (*right*) E_max_ for hβarr2, derived from data in *Panel j* and *Panel l*, respectively. Basal activity was evaluated by a *t*-test. All kinetic data represent the maximal response at each time point (Δ net BRET relative to baseline) and were fit by Michalis-Menton non-linear regression to derive K_m_ and V_max_. All E_max_ data were normalized to the WT-E_max_ for each respective agonist (set at 100%), then analyzed by two-way ANOVA followed by Tukey’s post-hoc test. **(o)** Dyn^K44A^ inhibiting receptor endocytosis. **(p)** Basal, **(q)** hGhrelin (C/R)-induced, **(r)** hGhrelin E_max_ for hβarr1 (derived from *Panel q*) to WT (grey) or L149P (light teal) co-expressing pcDNA or Dyn^K44A^. **(s)** Basal, **(t)** hGhrelin (C/R)-induced, and **(u)** hGhrelin E_max_ for hβarr2 (derived from *Panel t*) to WT (black/dark grey) or L149P (dark teal) co-expressing pcDNA or Dyn^K44A^. **(v)** MβCD-mediated depletion of membrane cholesterol (red) and βarr (light orange) recruitment to GHSR (blue). **(w)** Basal, **(x)** hGhrelin (C/R)-induced, and **(y)** hGhrelin E_max_ for hβarr1 (derived from *Panel* x) to WT (grey) or L149P (light teal) pretreated with 3mM MβCD (30-minutes, RT) or vehicle. **(a)** Basal, **(aa)** hGhrelin-induced, and **(bb)** hGhrelin E_max_ for hβarr2 (derived from *Panel aa*) to WT (black) or L149P (dark teal) pretreated with MβCD or vehicle. All basal and E_max_ data between *Panel*s *p-bb* were analyzed by two-way ANOVA followed by Tukey’s post-hoc test. All data represent the pooled mean ± SEM from ≥3 independent experiments in triplicate.

Next, we tested basal and agonist-stimulated βarr2 (**Figure 2h**) recruitment to the L149P and WT receptors using a similar BRET-based approach. In contrast to βarr1, basal βarr2 recruitment to the L149P mutant was reduced relative to the WT receptor (**Figure 2i**); whereas, ghrelin-induced βarr2 recruitment to L149P was equipotent to WT and efficacy was reduced (**Figure 2j**). Notably, the magnitude of ghrelin efficacy reduction for L149P-βarr2 (34%) was less than L149P-βarr1 (53%; see **Figure 2c**), indicating that the L149P mutant recruits βarr2 more effectively than βarr1, like the WT-GHSR ^14^. Furthermore, ghrelin-stimulated βarr2 association rate was similar between the L149P and WT receptors (**Figure 2i**). MK0677 potency for βarr2 recruitment at L149P was reduced mildly relative to the WT receptor, while MK0677 efficacy was reduced by only 24% at L149P (**Figure 2l**). Like ghrelin, MK0677-induced βarr2 recruitment association rate kinetics were similar between the WT and L149P receptors (**Figure 2m**). An analysis of ghrelin- and MK0677-βarr2 E_max_ showed that the L149P mutant had reduced agonist efficacy relative to the WT receptor (**Figure 2n**), albeit to a lesser extent than the reductions observed for βarr1 (see **Figures 2e, 2g**); moreover, MK0677-stimulated βarr2 recruitment efficacy to L149P was moderately increased relative to ghrelin efficacy (**Figure 2n**).

Recent work shows that agonist-directed endocytosis kinetics of Gα_q_-coupled GPCRs are predictive of βarr efficacy ^31^. To test this for L149P, we next co-expressed Dyn^K44A^ (**Figure 2o**) and measured βarr1/2 recruitment. Because MK0677 displayed a similar βarr1/2 recruitment profile to ghrelin, only ghrelin was tested in these experiments. Dyn^K44A^ co-expression did not affect basal βarr1 recruitment to the L149P or WT receptors (**Figure 2p**); whereas, Dyn^K44A^ markedly reduced ghrelin-stimulated βarr1 recruitment efficacy, but not potency, at both receptors (**Figure 2q-r**). Furthermore, Dyn^K44A^ co-expression significantly decreased ghrelin-βarr1 efficacy at the WT receptor to a level nearly identical to L149P in the absence of Dyn^K44A^ (**Figure 2q-r**). Basal βarr2 recruitment to L149P was unaffected by Dyn^K44A^ co-expression, while basal βarr2 recruitment to the WT receptor was reduced to a level comparable to L149P in the absence of Dyn^K44A^ (**Figure 2s**). Further, ghrelin-stimulated βarr2 recruitment to both L149P and WT was reduced upon Dyn^K44A^ co-expression, and the reduction observed for the WT receptor was comparable to L149P in the absence of Dyn^K44A^ (**Figures 2t-u**). These results together support that reduced basal βarr2 recruitment (see **Figure 2i**) and reduced agonist βarr1/2 efficacy (see **Figures 2c,g and 2j,n**) at the L149P mutant are likely related to its PM expression bias and/or distinct endosomal trafficking profile (see **Figure 1**). Furthermore, our findings support that the constitutive WT-GHSR endocytosis is required for basal βarr2, but not βarr1 recruitment; whereas, constitutive WT-GHSR endocytosis is required for the agonist-directed recruitment of both βarr1 and βarr2.

To test the effect of membrane cholesterol on βarr signaling, we next pretreated cells with MβCD (**Figure 2v**) or vehicle, then assessed basal and ghrelin-stimulated βarr1/2 recruitment to the L149P and WT receptors. MβCD pretreatment did not appreciably affect basal βarr1 recruitment to WT or L149P compared to their respective vehicle controls, but we did observe a mild reduction in basal βarr1 recruitment to the MβCD-pretreated L149P relative to the vehicle-pretreated WT receptor (**Figure 2w**). Ghrelin potency for βarr1 recruitment was moderately decreased at L149P (2.6-fold) and the WT (1.8-fold) receptor (**Figure 2x**). Furthermore, ghrelin E_max_ at L149P was significantly reduced by MβCD pretreatment, while a trend for reduced ghrelin E_max_ at the WT receptor did not reach statistical significance (**Figure 2y**). In contrast to βarr1, MβCD pretreatment decreased basal βarr2 recruitment to the WT receptor but did not affect L149P (**Figure 2z**), suggesting that reduced L149P-βarr2 recruitment is membrane cholesterol-dependent. Additionally, MβCD pretreatment attenuated ghrelin-induced βarr2 recruitment potency (**Figure 2aa**) and efficacy (**Figures 2aa-bb**) at both L149P and WT, although the effects were larger for the L149P mutant (**Figures 2aa-bb**) and similar in magnitude to βarr1 (see **Figures 2x-y**). Notably, MβCD pretreatment reduced ghrelin E_max_ at the WT receptor to a level statistically similar to the vehicle-pretreated L149P, suggesting that the moderate reduction in βarr2 recruitment efficacy to L149P at baseline (∼35%) is cholesterol-dependent. Together, these results support that (i) membrane cholesterol, at least partially, explains the selective reduction in basal βarr2 recruitment to the L149P mutant relative to βarr1 and that (ii) agonist-stimulated βarr1/2 recruitment to both the L149P and WT receptors is partially cholesterol-dependent, albeit more important for L149P.

Collectively, the results in **Figure 2** demonstrate that the L149P mutant recruits βarr1/2 with high agonist potency, moderately reduced efficacy, and normal association rate kinetics relative to the WT-GHSR. Additionally, L149P exhibits a selective reduction in basal βarr2 recruitment and higher agonist efficacy for βarr2 over βarr1, perhaps due to lower baseline recruitment. Our results also suggest that the βarr1/2 recruitment characteristics of L149P are, at least partially, related to its differential PM expression bias, endosomal trafficking, and association with membrane cholesterol (see **Figure 1**). For all descriptive data, C/R analyses, and statistics in **Figure 2**, see **Supplementary Table 2**.

### L149P-mediated inhibition of G protein signaling is Gα subtype-specific and cholesterol-independent

Next, we tested if the L149P mutant could couple to and/or activate Gα_q_—the preferential G protein for the GHSR ^6^. First, we employed the BRET-based TRUPATH assay to evaluate Gα_q_ activation, which measures Gα_q_-Gβγ dissociation upon guanine nucleotide exchange (GDP>GTP) ^32^ (**Figure 3a**). The GHSR exhibits remarkably high constitutive Gα_q_ activity (≥50% ghrelin E_max_ ^33^); thus, to provide a direct measure of constitutive activity, we included an empty vector (EV) condition to normalize basal receptor responses (*i.e.*, derive Δ net BRET relative to EV). Constitutive Gα_q_ dissociation from L149P was reduced by >70% relative to the WT receptor (**Figure 3b**). Furthermore, ghrelin- (**Figure 3c**) and MK0677- (**Figure 3d**) stimulated Gα_q_ dissociation was abolished at L149P, supporting that the L149P mutant confers complete βarr-bias (see **Figure 2**) relative to Gα_q_. Analysis of agonist E_max_ showed that both ghrelin and MK0677 agonism is abolished at L149P (**Figure 3e**), suggesting that the L149P mutation abrogates Gα_q_ activation by preferentially stabilizing an inactive conformation of the agonist-bound GHSR.

**Figure 3.**
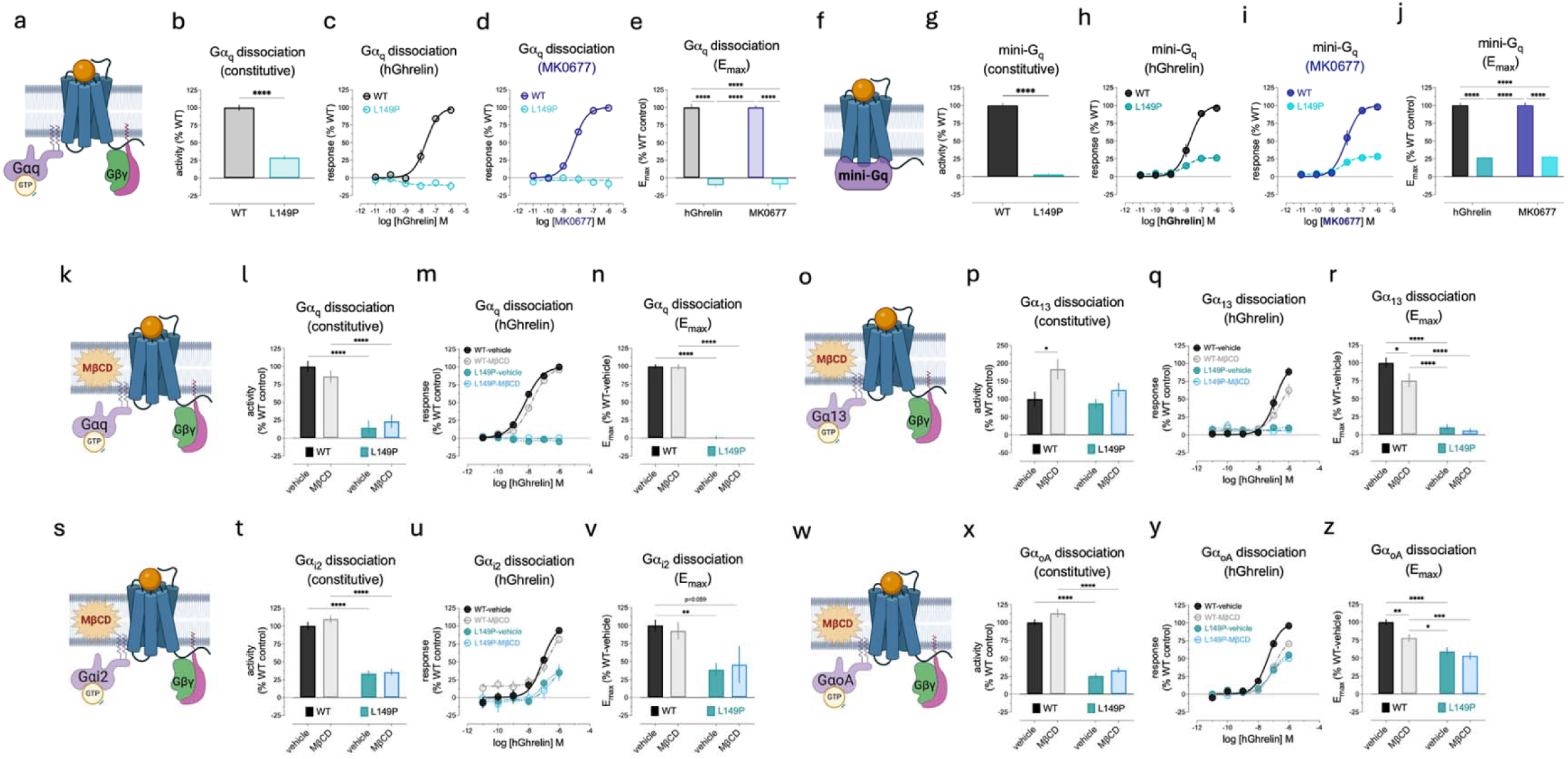
G protein signaling profile at the L149P mutant. **(a)** Gα_q_ dissociation into Gα_q_-GTP and free Gβγ, measured by TRUPATH. **(b)** Constitutive Gα_q_ activity of WT (grey) and L149P (light teal). **(c)** hGhrelin (C/R)-stimulated Gα_q_ dissociation at WT and L149P. **(d)** MK0677 (C/R)-stimulated Gα_q_ dissociation at WT (light blue) and L149P. **(e)** hGhrelin (*left*) and MK0677 (*right*) E_max_ for Gα_q_ dissociation, derived from data in *Panels c-d*. **(f)** mini-G_q_ coupling to GHSR (blue). **(g)** Constitutive mini-G_q_ recruitment of WT (black) and L149P (dark teal). **(h)** hGhrelin (C/R)-stimulated mini-G_q_ recruitment to WT and L149P. **(i)** MK0677 (C/R)-stimulated mini-G_q_ recruitment to WT (dark blue) and L149P. **(j)** hGhrelin (*left*) and MK0677 (*right*) E_max_ for mini-G_q_ recruitment, derived from data in *Panels h-i*. **(k)** Gα_q_ dissociation ± MβCD. **(l)** Constitutive, **(m)** hGhrelin (C/R), and **(n)** hGhrelin E_max_ for Gα_q_ at WT-vehicle (black), L149P-vehicle (dark teal), WT-MβCD (grey), and L149P-MβCD (light blue). **(o)** Gα_13_ dissociation ± MβCD. **(p)** Constitutive, **(q)** hGhrelin (C/R), and **(r)** hGhrelin E_max_ for Gα_13_. **(s)** Gα_i2_ dissociation ± MβCD. **(t)** Constitutive, **(u)** hGhrelin (C/R), and **(v)** hGhrelin E_max_ for Gα_i2_. **(w)** Gα_oA_ dissociation ± MβCD. **(x)** Constitutive, **(y)** hGhrelin (C/R), and **(z)** hGhrelin E_max_ for Gα_oA_. Constitutive activity was calculated by subtracting the absolute net BRET of EV/pcDNA plus biosensor (0%) from the absolute net BRET of receptor-alone to derive Δ net BRET, then normalizing to WT (100%). Constitutive activity was evaluated by a *t*-test (*Panels b* and *g*) or two-way ANOVA (*Panels l*, *p, t*, and *x*) followed by Tukey’s post-hoc test. E_max_ data were analyzed by two-way ANOVA followed by Tukey’s post-hoc test. All data represent the pooled mean ± SEM from ≥3 independent experiments run in duplicate or triplicate.

Subsequently, we tested if the L149P mutant could still couple to Gα_q_, despite its inability to activate the Gα_q_ heterotrimer via guanine nucleotide exchange. To test this, we employed the BRET-based mini-G_q_ sensor (**Figure 3f**), which mimics the nucleotide-free G protein active state and removes the Gα’s membrane anchoring, Gβγ binding, and α-helical domain ^34^. Constitutive mini-G_q_ recruitment was nearly abolished at the L149P mutant (**Figure 3g**); but to our surprise, ghrelin (**Figure 3h**) and MK0677 (**Figure 3i**) potency for mini-G_q_ recruitment at L149P was similar to the WT receptor, and agonist efficacy was reduced by ∼75% but not abolished (**Figure 3j**). Taken together with our TRUPATH results (see **Figures 3c-e**), these findings suggest that the L149P mutation selectively disrupts agonist-induced guanine nucleotide exchange and thus, may act as a guanine dissociation inhibitor (GDI), consistent with other ICL2 perturbations ^35, 36^.

The GHSR preferentially couples to Gα_q_, but it also couples with lower efficacy to G proteins within the Gα_i/o_ and Gα_12/13_ families—especially, Gα_i2_, Gα_oA_, and Gα_13_ ^6^. Thus, we next tested (i) if the L149P mutant activates these G proteins and (ii) if membrane cholesterol differentially affects signaling across G protein subfamilies. Using Gα_q_ as a control (**Figure 3k**), we confirmed both minimal L149P constitutive activity (**Figure 3l**) and no agonist-induced Gα_q_ activation (**Figures 3m-n**) under cholesterol-replete conditions. Following MβCD pretreatment, there was a mild, but statistically significant reduction in ghrelin-Gα_q_ potency at the WT, but not L149P mutant (**Figure 3m**), and no change in ghrelin efficacy at either receptor (**Figures 3l-n**). To our surprise, constitutive Gα_13_ (**Figure 3o**) activity was comparable between the vehicle-pretreated WT and L149P receptors; whereas, MβCD significantly increased WT constuitive activity and did not affect L149P (**Figure 3p**); whereas, ghrelin-stimulated Gα_13_ activation was abolished at L149P, while ghrelin efficacy was reduced at the WT receptor (**Figures 3q-r**). For Gα_i2_ (**Figure 3s**), both constitutive (**Figure 3t**) and ghrelin-stimulated activation (**Figures 3u-v**) were markedly reduced at L149P, but not abolished, in a cholesterol-independent manner for both receptors (**Figures 3t-v**). For Gα_oA_ (**Figure 3w**), constitutive (**Figure 3x**) and ghrelin-stimulated activation (**Figure 3y-z**) were markedly reduced, but not abolished, at L149P upon vehicle pretreatment; whereas, MβCD pretreatment decreased agonist potency and efficacy only at the WT receptor (**Figure 3y-z**).

In summary, the results in **Figure 3** support four key findings: The L149P mutant displays (i) nearly abolished Gα_q_ signaling, likely due to disrupted guanine nucleotide exchange [GDI activity]; (ii) no agonist-stimulated Gα_q_ or Gα_13_ activation; (iii) Gα_i/o_ partial agonism; (iv) normal Gα_13_ constitutive activity despite markedly reduced Gα_q_ and Gα_i/o_ constitutive activity; (v) cholesterol-independent effects on G proteins, while the WT-GHSR requires membrane cholesterol for most G protein signaling. For all descriptive data, C/R curve analyses, and statistics in **Figure 3**, see **Supplementary Table 4.**

### βarr recruitment to the L149P mutant requires GRK2/3-mediated phosphorylation but does not require G protein-dependent PKC or GRK2/3 activation

GHSR phosphorylation following Gα_q_^GTP^-dependent PKC activation and Gβγ-dependent GRK2/3 translocation ^37^ typically occurs upstream of βarr recruitment, with the latter mediated by Gβγ interaction with the pleckstrin homology (PH) domain of GRK2/3 ^38^. Accordingly, we next tested how the L149P mutant could elicit βarr-bias despite minimal G protein signaling (see **Figure 3**).

First, we used a pharmacological approach by employing a pan-PKC inhibitor, Go6983, or the selective GRK2/3 inhibitor, Compound 101 (Cmp101). To both test constitutive βarr recruitment and also control for potential confounding effects of pharmacological inhibitors (*e.g.*, due to Go6983 coloration), we included an EV control, as in our G protein assays (see **Figure 3**). For βarr1 (**Figure 4a**), Go6983-alone did not affect constitutive βarr1 recruitment to the WT receptor, while Cmp101-alone showed a trend for increased constitutive βarr1 recruitment but did not reach statistical significance. The combined pretreatment of Go6983-plus-Cmp101 reduced constitutive WT-βarr1 recruitment compared to Cmp101-alone but not vehicle (**Figure 4b**). In contrast, Go6983-alone moderately, and Cmp101-alone markedly, reduced ghrelin-stimulated WT-βarr1 recruitment (29% *versus* 83% decrease, respectively), while ghrelin potency was not affected (**Figures 4c-d**). Go6983-plus-Cmp101 had an additive effect, which nearly abolished WT-βarr1 recruitment efficacy (**Figures 4c-d**). At L149P, constitutive βarr1 recruitment was not affected by Go6983-alone, Cmp101-alone, or Go6983-plus-Cmp101 (**Figure 4e**). But in contrast to the WT receptor, Go6983-alone did not affect ghrelin-stimulated L149P-βarr1 recruitment efficacy; whereas, Cmp101-alone and Go6983-plus-Cmp101 reduced L149P-βarr1 recruitment efficacy to similar degrees (**Figure 4f-g**) and ghrelin potency was not affected (**Figure 4f**). For βarr2 (**Figure 4h**), similar effects were observed for both constitutive (**Figures 4i, 4l**) and agonist-stimulated βarr2 recruitment to the WT (**Figures 4j-k**) and L149P (**Figure 4m-n**) receptors, but with two exceptions: First, constitutive L149P-βarr2 recruitment following Cmp101-alone pretreatment trended towards an increase relative to vehicle but was only significantly different compared to Go6983-plus-Cmp101 (**Figure 4l**). Second, Go6983-plus-Cmp101 decreased ghrelin-stimulated L149P-βarr2 recruitment relative to Cmp101-alone. While unexpected, we speculate that this might reflect background PKC activity or signaling ‘crosstalk’ independent of L149P-mediated signaling. Overall, the results in **Figures 4a-n** support that the WT-GHSR requires both PKC and GRK2/3 for βarr1/2 recruitment, while L149P requires only GRK2/3.

**Figure 4.**
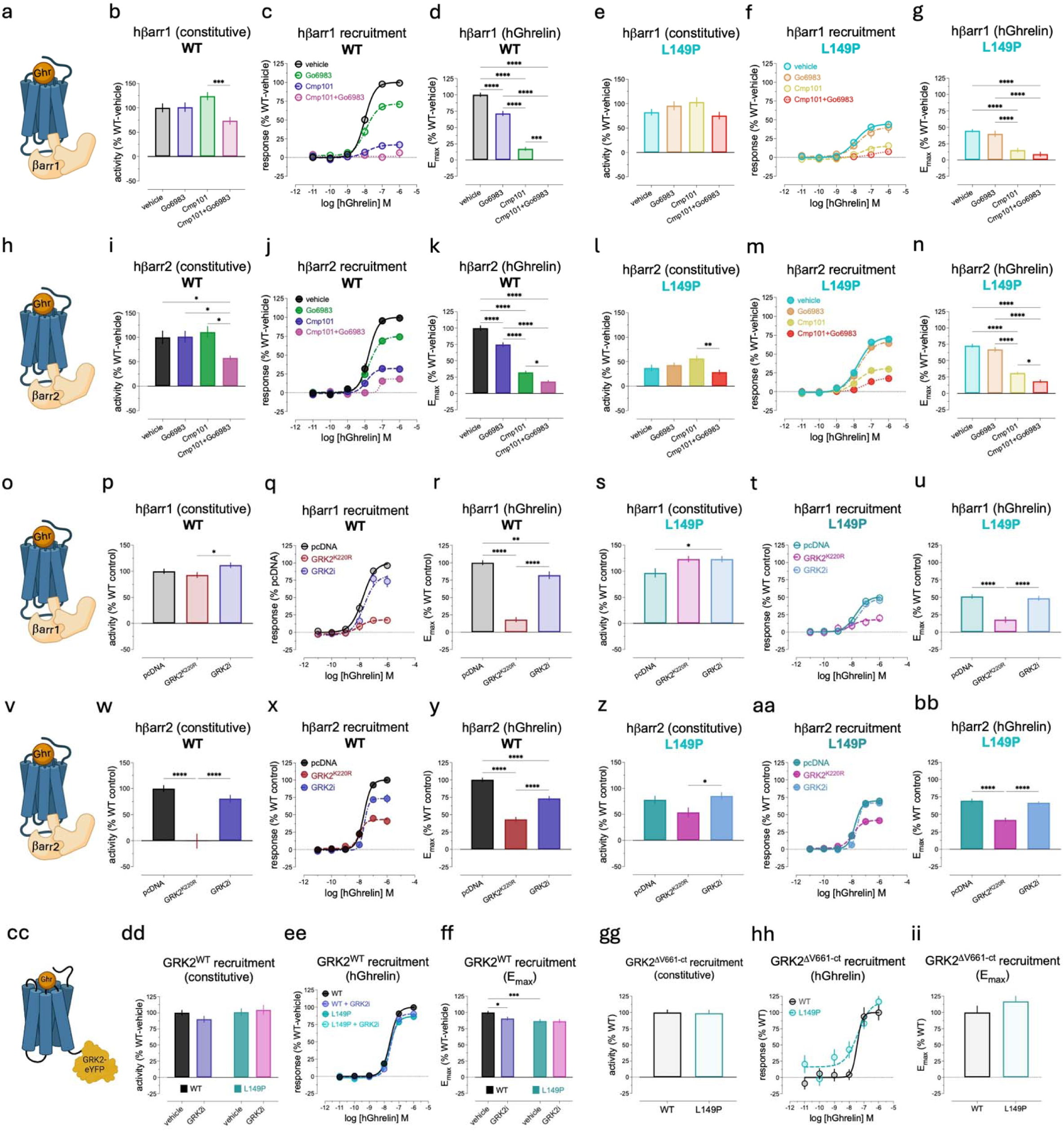
G protein-independent βarr1/2 and GRK2 recruitment to the L149P mutant. **(a)** hβarr1 recruitment to GHSR. **(b)** Constitutive, **(c)** hGhrelin C/R, and **(d)** hGhrelin E_max_ (derived from *Panel* c) of hβarr1 recruitment to WT following 30-minute pretreatment (RT) with vehicle (grey), Go6983 (light blue; 3μM), Cmp101 (light green; 10μM), or Go6983+Cmp101 (light purple). **(e)** Constitutive, **(f)** hGhrelin C/R, and **(g)** hGhrelin E_max_ (derived from *Panel f*) hβarr1 recruitment to L149P following pretreatment with vehicle (light teal), Go6983 (light orange), Cmp101 (light yellow), or Go6983+Cmp101 (light red). **(h)** hβarr2 recruitment to GHSR. **(i)** Constitutive, **(j)** hGhrelin C/R, and **(k)** hGhrelin E_max_ (derived from *Panel j*) of hβarr2 recruitment to WT following pretreatment with vehicle (black), Go6983 (dark blue), Cmp101 (dark green), or Go6983+Cmp101 (dark purple). **(l)** Constitutive, **(m)** hGhrelin C/R, and **(n)** hGhrelin E_max_ (derived from *Panel m*) hβarr2 recruitment to L149P following pretreatment with vehicle (dark teal), Go6983 (dark orange), Cmp101 (dark yellow), or Go6983+Cmp101 (dark red). **(o)** hβarr1 recruitment to GHSR. **(p)** Constitutive, **(q)** hGhrelin C/R, **(r)** hGhrelin E_max_ (derived from *Panel q*) of hβarr1 recruitment to WT following co-expression/pretreatment with pcDNA-vehicle (control; grey), GRK2^K220R^-vehicle (light red), or pcDNA-GRK2i (100μM, 30-minutes, RT). **(s)** Constitutive, **(t)** hGhrelin C/R, **(u)** hGhrelin E_max_ (derived from *Panel t*) of hβarr1 recruitment to L149P in pcDNA-vehicle (light teal), GRK2^K220R^-vehicle (light purple), or pcDNA-GRK2i (light blue). **(v)** hβarr2 recruitment to GHSR. **(w)** Constitutive, **(x)** hGhrelin C/R, **(y)** hGhrelin E_max_ (derived from *Panel x*) of hβarr2 recruitment to WT in pcDNA-vehicle (black), GRK2^K220R^-vehicle (dark red), or pcDNA-GRK2i (dark blue). **(z)** Constitutive, **(aa)** hGhrelin C/R, **(bb)** hGhrelin E_max_ (derived from *Panel aa*) of hβarr2 recruitment to L149P in pcDNA-vehicle (dark teal), GRK2^K220R^-vehicle (dark purple), or pcDNA-GRK2i (dark blue). All constitutive and E_max_ data were analyzed by one-way ANOVA. **(cc)** YFP-tagged WT-GRK2 (GRK2^WT^) recruitment to GHSR. **(dd)** Constitutive, **(ee)** hGhrelin C/R, **(ff)** hGhrelin E_max_ (derived from *Panel ee*) of GRK2^WT^ recruitment to WT-vehicle (black), WT-GRK2i (dark blue), L149P-vehicle (dark teal), and L149P-GRK2i (dark purple) following a 30-minute pretreatment (RT). Constitutive and E_max_ data were analyzed by two-way ANOVA followed by Tukey’s post-hoc test. **(gg)** Constitutive, **(hh)** hGhrelin C/R, **(ii)** hGhrelin E_max_ (derived from *Panel hh*) of recruitment of GRK2^ΔV661-ct^ to WT (grey) or L149P (light teal). Constitutive and E_max_ data were analyzed by *t*-test. All data represent the pooled mean ± SEM from ≥3 independent experiments in duplicate or triplicate.

To determine if GRK2/3-dependent βarr1 recruitment to the L149P mutant requires GRK2/3-mediated phosphorylation and/or Gβγ-dependent GRK2/3 translocation, we next co-expressed the phosphorylation-deficient GRK2 mutant, GRK2^K220R^, or pretreated cells with the Gβγ-GRK2/3 interfering peptide, GRK2i (**Figure 4o**). Constitutive βarr1 recruitment to the WT receptor was not affected by GRK2^K220R^ or GRK2i relative to control (pcDNA-vehicle). However, GRK2i mildly increased constitutive WT-βarr1 recruitment relative to GRK2^K220R^ (**Figure 4p**). Ghrelin-stimulated WT-βarr1 recruitment efficacy was markedly reduced by GRK2^K220R^ (82% decrease), and GRK2i moderately reduced ghrelin efficacy (19% decrease) (**Figure 4q-r**). Neither GRK2^K220R^ nor GRK2i affected ghrelin potency at the WT receptor (**Figure 4q**). At L149P, GRK2i mildly increased constitutive βarr1 recruitment relative to control (**Figure 4s**). Importantly, while GRK2^K220R^ markedly reduced L149P-βarr1 recruitment efficacy, GRK2i had no effect (**Figure 4t-u**). Ghrelin potency for L149P-βarr1 recruitment was not affected by GRK2^K220R^ or GRK2i (**Figure 4t-u**). For βarr2 (**Figure 4v**), GRK2^K220R^ abolished WT constitutive activity, while GRK2i had no effect (**Figure 4w**). Similar to βarr1, ghrelin-stimulated WT-βarr2 recruitment efficacy was markedly reduced by GRK2^K220R^ (57% decrease) and moderately reduced by GRK2i (28% decrease), while ghrelin potency was unaffected (**Figure 4x-y**). At L149P, there was a trend for reduced constitutive βarr2 recruitment upon GRK2^K220R^ co-expression, but this effect did not reach statistical significance relative to control. We did, however, identify a small statistically significant increase in constitutive L149P-βarr2 recruitment following GRK2i pretreatment compared to GRK2^K220R^ co-expression (**Figure 4z**). Similar to βarr1, GRK2^K220R^ reduced ghrelin-stimulated L149P-βarr2 recruitment efficacy and GRK2i had no effect, while ghrelin potency was not affected in either condition (**Figure 4aa-bb**). Together, these results support that L149P-βarr1/2 recruitment requires GRK2/3-mediated receptor phosphorylation but not Gβγ interaction; whereas, the WT-GHSR requires both of these processes for βarr1/2 recruitment.

To further test if L149P recruits GRK2/3 in a Gβγ-independent manner, we performed BRET-based receptor-GRK2^YFP^ recruitment assays ^28^ (**Figure 4cc**) in the presence or absence of GRK2i. Constitutive GRK2 recruitment was similar between WT and L149P under vehicle-treated conditions, and GRK2i did not affect constitutive GRK2 recruitment to either receptor (**Figure 4dd**). Consistent with our prediction, GRK2i pretreatment decreased WT-GRK2 recruitment efficacy, albeit mildly (10% decrease), and did not affect L149P-GRK2 recruitment efficacy (**Figure 4ee-ff**). Ghrelin potency was not affected by GRK2i pretreatment for either receptor (see **Figure 4ee**). We suspect that the moderate effect of GRK2i on GRK2 recruitment to the WT receptor was due to GRK2 over-expression in these experiments. Notably, the reduction WT-GRK2 recruitment following GRK2i pretreatment was statistically equivalent to L149P-GRK2 recruitment under vehicle-treated conditions (see **Figure 4ff**), supporting that reduced L149P-GRK2 recruitment at baseline is related to a lack of Gβγ-mediated GRK2 translocation. To further test this, we utilized a truncated GRK2 mutant, GRK2^ΔV661-ct^, which abrogates the Gβγ-GRK2 interaction by removing the far C-terminal Gβγ-binding (PH) domain of GRK2 ^28^. Consistent with our GRK2i results (see **Figure 4dd**), there was no difference in constitutive (**Figure 4gg**) or ghrelin-stimulated (**Figure 4hh-ii**) GRK2^ΔV661-ct^ recruitment between the WT and L149P receptors (see **Figure 4ee-ff**).

To test if Gα_i/o_ partial agonism (see **Figures 3s-z**) could explain Gβγ-independent βarr1/2 recruitment to L149P, we pretreated cells with Gα_i/o_ inhibitor—pertussis toxin (PTX) ^39^—then tested ghrelin-stimulated βarr1/2 recruitment. Ghrelin-induced βarr1 recruitment efficacy, but not potency, at the WT receptor was reduced moderately by PTX (**Supplementary Figure 2a**), while L149P-βarr1 recruitment was unaffected (**Supplementary Figures 2b-c**). For βarr2, ghrelin potency and efficacy were not affected by PTX at either receptor (**Supplementary Figure 2d-f**). Thus, Gα_i/o_ signaling does not likely account for Gβγ-independent βarr1/2 (see **Figures 4aa-bb**) or GRK2 recruitment (see **Figures 3ee-ff**, **4hh-ii**) to L149P.

Overall, the results in **Figure 4** support that, in contrast to the WT-GHSR, the L149P mutant recruits βarr1/2 and GRK2 without G protein engagement, independently of PKC, Gβγ-dependent GRK2/3 translocation, and Gα_i/o_. Nevertheless, the L149P mutant still requires enzymatic activation of GRK2/3 (receptor phosphorylation) for βarr1/2 recruitment. For all descriptive data, C/R curve analyses, and statistics in **Figure 4**, see **Supplementary Table 5**.

### The L149P mutation switches the GRK recruitment profile of the GHSR

βarr-biased ligands and receptors preferentially utilize GRK5/6 over GRK2/3 for βarr recruitment ^26, 27^. Thus, we next tested if the L149P mutation affects the receptor’s GRK preference profile. To this end, we tested βarr1/2 recruitment to the WT and L149P receptors in HEK293/A cell lines lacking all GRKs (ΔGRK2/3/5/6), GRK2/3 (ΔGRK2/3), or GRK5/6 (ΔGRK5/6) ^26^.

For βarr1 (**Figure 5a**), WT constitutive activity was not affected by loss of any GRK subclasses (**Figure 5b**). Conversely, ghrelin-stimulated WT-βarr1 recruitment efficacy was abolished by ΔGRK2/3/5/6 and ΔGRK2/3, while ghrelin potency and efficacy were not affected by ΔGRK5/6 (**Figures 5c-d**), supporting that agonist-induced WT-βarr1 recruitment is entirely GRK2/3-dependent. Constitutive L149P-βarr1 recruitment was also not affected by loss of any GRK subclasses (**Figure 5e**). Ghrelin-stimulated L149P-βarr1 recruitment efficacy was abolished in ΔGRK2/3/5/6 cells; however, while trends for decreased L149P-βarr1 recruitment were observed in ΔGRK2/3 and ΔGRK5/6 cells (**Figures 5f-g**), the effect did not reach significance for ΔGRK5/6 and statistical comparisons were precluded for ΔGRK2/3 due to high background and poor curve fit. For βarr2 (**Figure 5h**), constitutive recruitment to the WT receptor was not affected by knockout of any GRK subclasses (**Figure 5i**); whereas, ghrelin-stimulated WT-βarr2 recruitment efficacy was abolished by ΔGRK2/3/5/6, markedly reduced by ΔGRK2/3, and increased by ΔGRK5/6, further supporting that the WT-GHSR requires only the GRK2/3 subclass for βarr recruitment (**Figures 5j-k**). We speculate that the increase in WT-βarr2 recruitment in ΔGRK5/6 cells is due to a compensatory upregulation of GRK2/3, as observed elsewhere ^26^. Ghrelin potency was not affected in any conditions for the WT receptor (**Figure 5j**). For L149P, constitutive βarr2 recruitment was not affected by total GRK knockout (**Figure 5l**); whereas, ghrelin-stimulated L149P−βarr2 recruitment efficacy was markedly reduced in ΔGRK2/3/5/6 and ΔGRK2/3 cells (**Figures 5m-n**), consistent with our results using pharmacological inhibitors (see **Figure 4m-n**). But in contrast to the WT-GHSR, ghrelin-stimulated βarr2 recruitment to L149P was reduced in ΔGRK5/6 cells (**Figures 5m-n**). Ghrelin potency at L149P was not affected by knockout of any GRK subclasses (**Figure 5m**).

**Figure 5.**
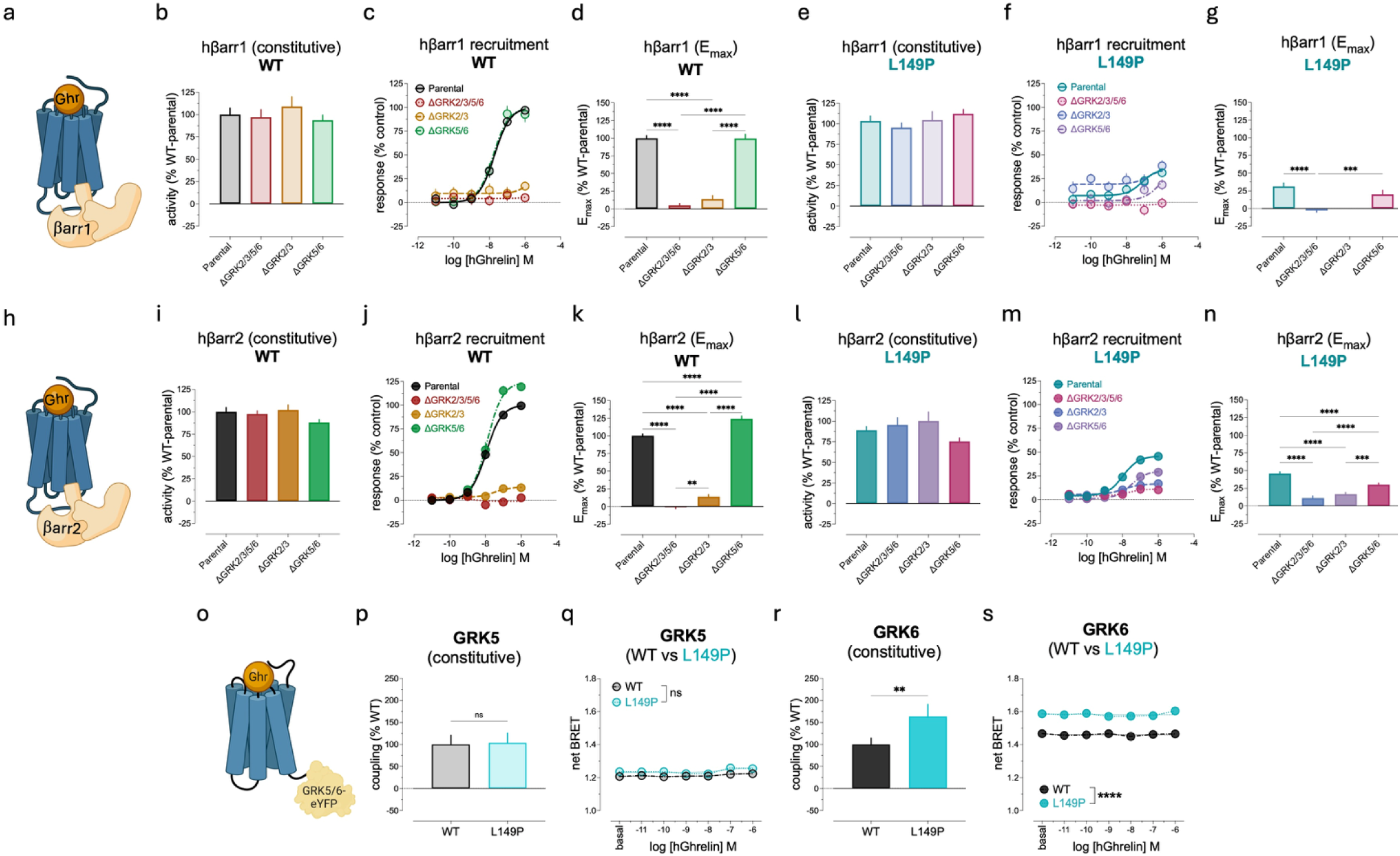
GRK subtype-specific βarr1/2 recruitment and GRK5/6 coupling to the L149P mutant. **(a)** hβarr1 recruitment to GHSR. **(b)** Constitutive, **(c)** hGhrelin-stimulated C/R, and **(d)** hGhrelin E_max_ (derived from *Panel c*) of hβarr1 recruitment to WT in parental (light grey), ΔGRK2/3/5/6 (light red), ΔGRK2/3 (light orange), and ΔGRK5/6 (light green). **(e)** Constitutive, **(f)** hGhrelin-stimulated C/R, and **(g)** hGhrelin E_max_ (derived from *Panel f*) of hβarr1 recruitment to L149P in parental (light teal), ΔGRK2/3/5/6 (light blue), ΔGRK2/3 (light purple), and ΔGRK5/6 (light maroon). **(h)** hβarr2 recruitment to GHSR. **(i)** Constitutive, **(j)** hGhrelin-stimulated C/R, and **(k)** hGhrelin E_max_ (derived from *Panel j*) of hβarr2 recruitment to WT in parental (dark grey), ΔGRK2/3/5/6 (dark red), ΔGRK2/3 (dark orange), and ΔGRK5/6 (dark green). **(l)** Constitutive, **(m)** hGhrelin-stimulated C/R, and **(n)** hGhrelin E_max_ (derived from *Panel m*) of hβarr2 recruitment to L149P in parental (dark teal), ΔGRK2/3/5/6 (dark blue), ΔGRK2/3 (dark purple), and ΔGRK5/6 (dark maroon). **(o)** GRK5/6^YFP^ coupling to GHSR. **(p)** Constitutive and **(q)** hGhrelin C/R-treated GRK5^YFP^ coupling, then **(r)** constitutive and **(s)** hGhrelin C/R-treated GRK6^YFP^ coupling over 30-minutes. *Panel p*, *r* were analyzed by Mann-Whitney nonparametric test; whereas, *Panels q, s* were analyzed by a mixed-model two-way ANOVA. All data represent the pooled mean ± SEM from 4 independent experiments in triplicate.

To test GRK5/6 coupling directly, we employed a BRET-based receptor-GRK5/6^YFP^ recruitment assay (**Figure 5o**). While constitutive (**Figure 5p**) and ghrelin-stimulated (**Figures 5q**) GRK5 coupling to L149P did not differ from the WT receptor, constitutive L149P-GRK6 coupling was markedly increased (**Figure 5r**) and remained elevated following ghrelin treatment (**Figures 5s**). Ghrelin stimulation did not increase GRK5/6 coupling at either receptor relative to their respective baselines (**Figures 5q, 5s**); thus, we expressed the data as non-normalized, absolute net BRET (**Figures 5p-s**).

Collectively, the results in **Figure 5** support that agonist-induced βarr recruitment to the L149P mutant is GRK2/3 and GRK5/6-dependent, while βarr recruitment to the WT-GHSR is exclusively GRK2/3-dependent under endogenous GRK expression. Furthermore, the L149P mutation markedly increases constitutive GRK6 coupling and maintains GRK6 coupling following agonist treatment; thus, selective GRK5/6-dependent βarr recruitment to L149P might be due to enhanced GRK6 ‘precoupling’. For all descriptive data, C/R analyses, and statistics in **Figure 5**, see **Supplementary Table 6**.

## DISCUSSION

In this study, we identify and characterize an ultra-rare, coding variant in Leu149^ICL2^ [34.51] of the GHSR—a prototypical rhodopsin-like GPCR. Our group has shown previously that the Leu149^ICL2^ residue, proximal to the highly conserved E/DRY motif, regulates transducer coupling ^10^ and receptor trafficking ^14^. Herein, we expand upon this work by demonstrating that a natural missense mutation in Leu149^ICL2^ induces conformational states unfavorable to cholesterol-enriched PM microdomains and G protein activation, despite robust βarr1/2 recruitment and agonist-directed endosomal translocation. Furthermore, the L149P mutant switches the GRK selectivity profile of the GHSR from an exclusively GRK2/3-dependent, to a GRK2/3/5/6-dependent GPCR, which recruits βarr1/2 in a G protein-independent manner. Collectively, our results highlight how a single, naturally-occurring substitution in ICL2 can simultaneously influence receptor–lipid interactions, biased signaling, and trafficking.

First, we found that the natural L149P mutant enhances basal PM localization via decreased constitutive internalization/cycling and distrupts receptor distribution within its local lipid microenvironment, including diminished clustering within cholesterol-enriched lipid rafts and reduced lipid raft-mediated endocytosis. These effects may reflect changes in ICL2 conformation and/or mobility, as the orientation of GPCR transmembrane helices and loops across inactive-to-active receptor states determine their hydrophobic thickness, thereby impacting lipid interactions and partitioning ^29^. The L149P mutant preferentially localized within putative non-raft PM microdomains, typically comprised of more disordered unsaturated lipids, suggesting that ICL2 perturbations might favor partioning into more fluid membrane compartments by imparting increased receptor flexibility. The GHSR regulates systemic energy balance via dynamic nutrient sensing and is sensitive to diet-derived nutrients ^40^; thus, dietary changes in lipid composition that alter the PM microenvironment could affect GHSR bias and trafficking.

Additionally, ghrelin-stimulated L149P endosomal translocation was enhanced, suggesting that perturbation of ICL2 may also redirect intracellular trafficking routes. As a Class A GPCR, the WT-GHSR exhibits rapid, agonist-induced βarr recruitment and minimal endosomal translocation. In contrast, the L149P mutant displayed robust agonist-directed endosomal translocation in a Class B-like manner. Given divergent GRK subtype dependence between the WT and L149P receptors, and enhanced L149P-GRK6 precoupling, these results suggest that distinct ICL3 or C-tail phosphorylation patterns (‘barcodes’) could account for distinct trafficking of the L149P mutant. Moreover, considering similar total internalization of the WT and L149P receptors determined by ELISA, yet slower WT-GHSR endosomal translocation kinetics and lipid raft enrichment, it is possible that the WT-GHSR internalizes, in part, via caveolin-dependent pathways while L149P preferentially utilizes clathrin-mediated endocytosis.

Based on prior work demonstrating that ICL2 forms a short α-helix in active GPCR–transducer complexes ^41, 42^, we speculate that “helix-breaking” Leu149^ICL2^ mutations might reduce ICL2 α-helical stability and disrupt ICL2-E/DRY motif interactions critical to G protein activation. Despite a lack of Gα_q_ activation, the L149P mutant elicits high-affinity/low-efficacy mini-G_q_ coupling upon agonist stimulation, supporting prior work showing that GPCR^ICL2^ mutations can prevent guanine nucleotide exchange without precluding G protein coupling ^43^. Furthermore, the L149P mutant exhibits partial agonism at Gα_i/o_, while agonist-induced Gα_q_ and Gα_13_ signaling were abolished, consistent with Gα_i_ requiring less ICL2 α-helical stability relative to other G proteins ^44^. Importantly, membrane cholesterol is required partially for both constitutive and agonist-stimulated G protein signaling at the WT-GHSR, while any observable L149P-mediated G protein signaling is cholesterol-independent. Moreover, cholesterol depletion alters WT receptor-mediated G protein signaling in a manner directionally consistent with L149P under cholesterol-replete conditions (*e.g.*, MβCD-induced increase in constitutive WT-Gα_13_ activity and attenuated agonist-induced Gα_q_, Gα_13_, and Gα_oA_ activation). Critically, these results contrast with βarr, wherein cholesterol depletion disproportionately affects the L149P mutant. Given the differential distribution of Gα_q_, Gα_13_, and Gα_i/o_ in caveolae ^45^, non-rafts ^46^, and lipid rafts ^45^, and the preferential recruitment of βarr to non-rafts ^47^, L149P-βarr bias could be related, in part, to PM partitioning-dependent transducer proximity.

Despite markedly reduced G protein activation, the L149P mutant retains high intrinsic efficacy for βarr via mechanisms dependent partially on both its PM expression bias and membrane cholesterol. These results support well-established ideas suggesting that (i) ICL2 conformations critically regulate transducer coupling selectivity, (ii) G protein- and βarr-coupled GPCRs can adopt distinct conformational states, and (iii) βarr recruitment is dependent on receptor subcellular distribution and microdomain partitioning [location bias]. Furthermore, L149P recruits βarr independently of both PKC and Gβγ-dependent GRK2/3 translocation, supporting that βarr recruitment can, in some cases, be dissociated from G protein signaling. Lastly, we found that the L149P mutant requires both endogenous GRK2/3 and GRK5/6 for βarr recruitment, while the WT-GHSR requires only endogenous GRK2/3. These results are consistent with other βarr-biased ligands ^26^ and receptors ^27^ that preferentially utilize GRK5/6. Because of their membrane-anchoring properties, the L149P mutant may disproportionately recruit GRK5/6 due to its PM expression bias and/or differential PM partitioning. Consistent with this, the L149P mutant exhibits elevated constitutive GRK6 association and sustained GRK6 proximity following agonist stimulation. Thus, enhanced GRK6 pre-coupling could ‘prime’ L149P conformations for βarr coupling, desensitization, and Class B-like receptor trafficking. Broadly, we propose that the Leu149^ICL2^ locus is a conserved “bias hotspot” and thereby provides a structural blueprint to rationally design βarr–biased ligands. Harnessing this “bias hotspot” pharmacologically with biased allosteric modulators may offer an advantageous approach to achieve therapeutic efficacy while simultaneously reducing side effects ^48^.

Although the clinical relevance of this ultra-rare *GHSR* SNP (rs1390872592) is currently unknown, our findings underscore how single amino acid mutations can confer dramatic changes to GPCR structure/function. In a physiological context, we speculate that the L149P mutation might reduce energy intake, body weight, and growth hormone secretion, as well as improve glucose control, by reducing GHSR-mediated Gα_q_ signaling ^9^. Because the ICL2^34^.^51^ site and neighboring residues (*e.g.*, GHSR-Pro148^34^.^50^) are well-conserved across rhodopsin-like GPCRs, it is conceivable that other rare variants in this locus could produce pathophysiological effects. In our companion paper (Zhou *et al.*), we show that rare natural variants in the corresponding hydrophobic ICL2 locus^34^.^50–34.51^ of phylogenetically-related neurotensin-1 receptor (NTSR1) engender biased signaling phenotypes similar to GHSR-L149P. Thus, we propose that the hydrophobic ICL2^34.50–34.51^ motif is highly conserved due to its important regulation of ICL2 conformational state dynamics and transducer coupling. Consistent with this, a study published during the writing of this manuscript found that β_2_AR biased signaling requires Phe139^34.51^ via regulation of ICL2 α-helical stability ^44^, further supporting our findings herein. Thus, while natural mutations in the hydrophobic ICL2 locus^34.50–34.51^ locus are rare, their importance in receptor signaling, subcellular distribution, and trafficking could plausibly manifest clinically-relevant effects when extrapolated across large, heterogenous populations.

**Figure 6.**
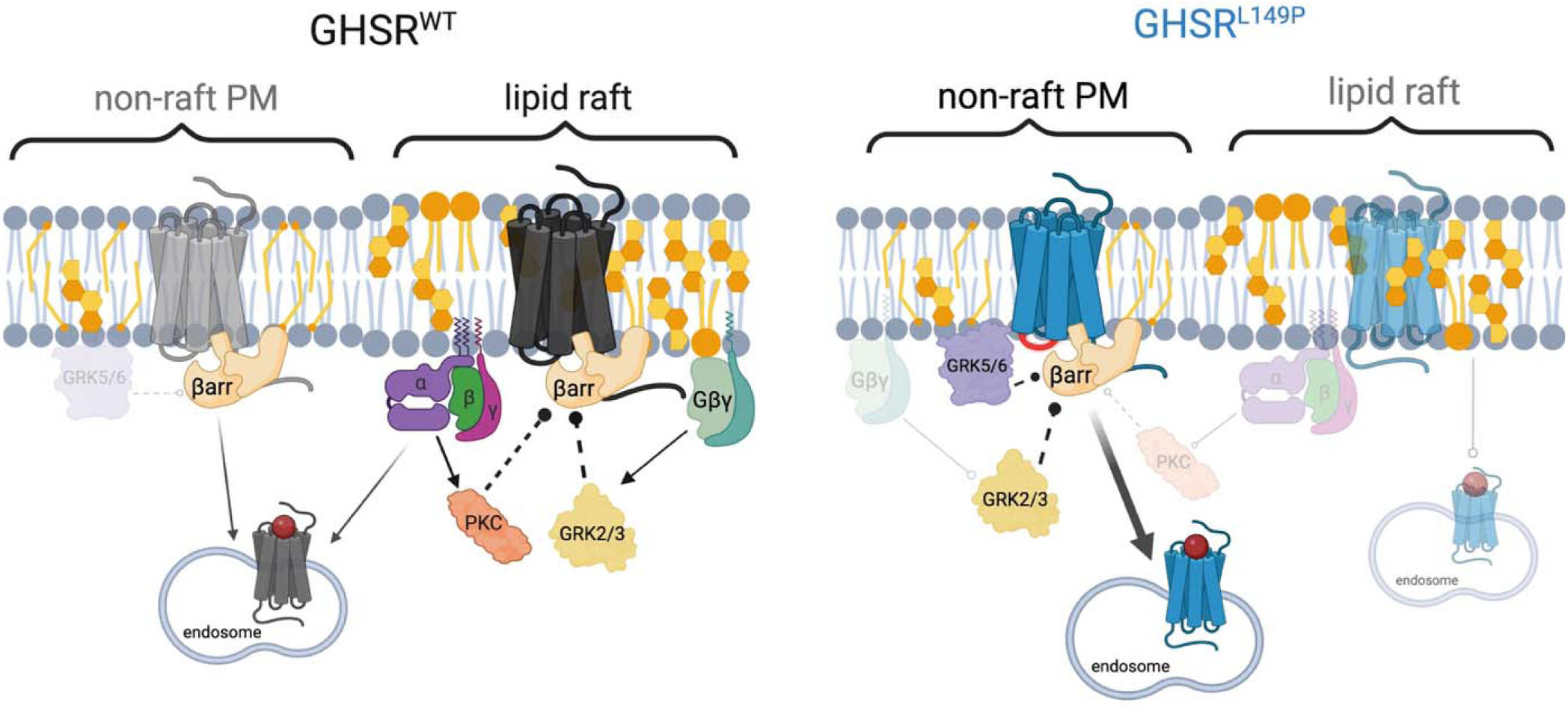
Simplified model of hypothesized L149P mechanisms. We hypothesize that the ultra-rare, natural L149P mutant (teal, *right*), destabilizes the α-helical conformation of ICL2, thereby (i) reducing clustering within cholesterol/sphingolipid-enriched lipid rafts and (ii) blocking G protein activation. Consequently, L149P preferentially partitions into more disordered, non-lipid raft PM microdomains (*e.g.*, unsaturated fatty acid-enriched) and exhibits poor agonist-directed endocytosis from lipid rafts. Under basal conditions, L149P expression is biased towards the PM over endosomes due to attenuated constitutive internalization/receptor cycling. However, upon agonist stimulation, L149P undergoes robust internalization and endosomal translocation via Class B-like kinetics distinct from the WT-GHSR (black, *left*), suggesting that L149P may utilize different phosphorylation ‘barcodes’ and/or endocytic routes. The L149P mutant readily recruits βarr in a partially cholesterol-dependent manner, while simultaneously inhibiting Gα_q_ via decreased guanine nucleotide exchange. Accordingly, L149P does not activate PKC or Gβγ-dependent GRK2 translocation and recruits βarr in G protein-independent manner despite weak Gα_i/o_ partial agonism. In contrast to the WT-GHSR, βarr recruitment to L149P requires both endogenous GRK2/3 and GRK5/6, with the latter associated with increased L149P-GRK6 preassociation. Thus, GRK6 precoupling to L149P might ‘prime’ the receptor towards a constitutively βarr-favoring conformational state readily amendable to βarr coupling, desensitization, and endocytosis.

## MATERIALS & METHODS

### Cell Culture & Transfections

HEK293/T-17 cells (Cat no. CRL-11268) were purchased from the American Type Culture Collection (ATCC). HEK293/A parental (ThermoFisher; Waltham, MA) and their derivative CRISPR/Cas9-engineered GRK knockout (DGRK2/3/5/6, DGRK2/3, DGRK5/6) were described previously ^26^. All cell lines were maintained in Dulbecco’s Modified Eagle Medium (DMEM; Gibco, ThermoFisher; Cat no. 11995073) supplemented with 10% fetal bovine serum (FBS; Gibco, ThermoFisher; Cat no. 26140095) and antibiotic-antimycotic (Gibco, ThermoFisher; Cat no. 15240062) (complete DMEM). Cells were cultured in a humidified 5% CO_2_ incubator at 37°C and grown on coated tissue culture plates. All cell lines were routinely split/passaged using trypsin/EDTA (Gibco, ThermoFisher; Cat no. 25200114) and never exceeded >30 passages.

For all transfections, cells were first seeded at a density of 6-7.5 x 10^5^ cells/well in a 6-well plate. Then 16-24 hours later, cells were transfected using a standard calcium phosphate method. Transfection reagents — CaCl_2_ (2.5M; Cat no. TBS5072) and HEPES-buffered saline (2xHBS, pH 7.05; Cat no. TBS5076) — were purchased from TriBioscience (Sunnyvale, CA). As described above, L149P was transfected at a 0.5:1 ratio relative to the WT-GHSR (typically 50ng:100ng, respectively) to adjust for total receptor expression (see **Figure 1c**), except when indicated otherwise (see **Figure 1b**). Dyn^K44A^, ^mVenus^hβarr1/2, and all GRK^YFP^ plasmids were transfected at a mass of 1mg, while TRUPATH sensors were all transfected at a mass of 100ng. Filler cDNA (pcDNA3.1+ or pcDNA5 for TRUPATH experiments) was used to bring the total transfection reaction mixture up to 2-3mg. Six-eight hours after transfection, the media was exchanged for complete DMEM, then incubated overnight prior to 96-well plate seeding (described below). All cell counting and transfection efficiency measurements were performed using a Countess 3 FL automated cell counter (ThermoFisher; Cat no. AMQAF2000).

### Plasmids & Pharmacological Reagents

Plasmid cDNA synthesis and site-directed mutagenesis of the N-terminally hemagglutinin (3xHA)-tagged hGHSR_1a_^WT^, hGHSR_1a_^L149P^, hGHSR_1a_^WT^-RLuc8, hGHSR_1a_^L149P^-RLuc8 was performed by GenScript (Piscataway, NJ) and cloned into a pcDNA3.1+ vector backbone. For empty vector control conditions to measure constitutive activity (see **Figures 3-5**), pcDNA3.1+ was used. The rat dynamin-1 dominant negative, Dyn^K44A^, was generated by the Caron Laboratory (Duke University; Durham, NC) and described previously ^49^. The bBRET sensors mVenus-CAAX ^50^, (MyrPalm)-mVenus ^6^, and 2xFYVE-mVenus ^51^ have been described previously. The N-terminally mVenus-tagged human βarr1 (hβarr1) and βarr2 (hβarr2) plasmids were a gift from Dr. Lauren Slosky (University of Minnesota; Minneapolis, MN). All TRUPATH sensors were gifts from Dr. Bryan Roth (Addgene Kit #1000000163). The mini-G_q_ plasmid was a gift from Dr. Nevin Lambert (Augusta University; Augusta, GA). The dominant negative mutant, bovine GRK2-K220R, was a gift from Dr. Robert Lefkowitz (Addgene plasmid #35403). The GRK2^WT^-YFP ^28^, GRK2^V661-ct^-YFP ^28^, and GRK5/6-YFP ^52^ plasmids were described previously.

Human acyl-ghrelin (ghrelin), MK0677, Go6983, and Cmp101 were purchased from Tocris Bioscience (Bristol, UK). MβCD and GRK2i were purchased from MedChemExpress (Monmouth Junction, NJ). Pertussis toxin (PTX) was purchased from Cayman Chemical (Ann Arbor, MI).

### On- and In-Cell ELISA Assays

ELISA assays were performed as previously described ^6^. Briefly, transfected cells were seeded on poly-D-lysine-coated 96-well plates at a density of ∼45,000 cells/well in phenol red-free opti-MEM (Gibco, ThermoFisher; Cat no. 11058021) supplemented with 2% FBS and antibiotic-antimycotic (‘BRET media’). After overnight culture, BRET media was aspirated and serum-starved in opti-MEM (containing no FBS or antibiotic-antimycotic) for 2-4 hours prior to stimulation with vehicle (opti-MEM), ghrelin (1μM), or MK0677 (1μM) for 45-minutes at 37°C. Next, media was rapidly aspirated and fixed in 4% paraformaldehyde for 15-minutes at RT. Fixed cells were washed three times (∼5-minutes/wash) in phosphate-buffered saline (PBS, pH 7.4; Gibco), then blocked for 1-hour in fish gelatin (Rockland Immunochemicals; Philadelphia, PA; Cat no. MB-067-0100). Cells were then probed with horseradish peroxidase (HRP)-conjugated anti-HA polyclonal antibody (Invitrogen, Cat no. PA1-29751) for 1-hour at RT (1:2,500). Next, cells were washed three times with PBS, then incubated with 1x SuperSignal^TM^ West Pico PLUS Chemiluminescent Substrate (ThermoFisher; Cat no. 34577) for ∼5-minutes prior to measuring luminescence in a TriStar3 multimodal plate reader (Berthold Technologies; Baden-Württemberg, Germany). For in-cell ELISA assays, blocking buffer, antibody buffer, and PBS wash buffer contained 0.2% triton-X-100 to permeabilize the cells. For on-cell ELISA assays, triton-X-100 was omitted in order to label only surface receptors.

### bBRET and BRET1/2 Assays

All bBRET and BRET1/BRET2 assays were performed as previously described ^6^. Briefly, transfected cells were seeded on poly-D-lysine-coated 96-well plates at a density of 40-45,000 cells/well in BRET media. After overnight culture, cells were serum-starved in Hank’s Balance Salt Solution (HBSS; Gibco; Cat no. 14025134) supplemented with 20mM HEPES for 2-4 hours at 37°C. Then, cells were either pretreated with inhibitors for 30-minutes at RT or proceeded immediately to agonist treatment. Five-ten minutes prior to agonist treatment, cells were incubated with coelenterazine-H (NanoLight Technology; Pinetop, AZ; Cat no. 301) for BRET1 (mVenus/eYFP) or coelenterazine-400a (NanoLight; Cat no. 340) for BRET2 (GFP2) based on manufacturer instructions. All BRET measurements were taken on a TriStar3 plate reader set to 37°C for bBRET assays or RT (22-25°C) for all other assays using the recommended filter settings. All ‘*basal*’ activity measurements represent the maximal absolute net BRET (acceptor/donor) of receptor-alone conditions (*i.e.*, non-normalized); whereas, all *‘constitutive*’ activity measurements represent the maximal difference between net BRET of receptor-alone minus the net BRET of pcDNA/EV-alone (*i.e.*, to derive Δ net BRET). All agonist-stimulated response data represent the maximal difference in the net BRET of agonist-stimulated conditions (at each ligand concentration) minus the net BRET of receptor-alone (*i.e.*, agonist Δ net BRET). To control for time-dependent effects, all Δ net BRET measurements (constitutive and agonist) were calculated/normalized relative to its respective control within the same column on a 96-well plate.

### Statistics & Data Analysis

As indicated in the Figure Legends, all data are presented as the pooled mean±SEM derived from ≥3 independent experiments. For all biochemical experiments, 2-4 technical replicates were included. All data were plotted and analyzed in GraphPad Prism version 10 with a statistical significance threshold defined as p<0.05; ns, p>0.05, *, p<0.05; **, p<0.01; ***, p<0.001; ****, p<0.0001. All data represent the maximal responses over the full assay duration following vehicle or agonist treatment and are normalized to the WT top/E_max_ (100%) and bottom (0%), then fit by a three- or four-parameter non-linear regression determined statistically by an F-test.

## AUTHOR CONTRIBUTIONS

*Conceptualization*: A.T., L.S.B. and J.D.G.; *Methodology*: A.I. and J.D.G.; *Investigation/Performed Experiments*: E.M.B., L.K., J.A.S., and J.D.G.; *Data/Statistical Analysis*: A.T. and J.D.G.; *Writing*: A.T. and J.D.G.; *Review/Editing*: E.M.B., A.T., L.K., A.I., L.S.B., and J.D.G.

## DATA & MATERIALS

All data required to evaluate the results and conclusions in this manuscript are presented in the main and supplementary figures and tables. All materials generated for, or used in, this manuscript are described in the *Material & Methods*.

## REFERENCES

1. Hauser AS, Attwood MM, Rask-Andersen M, Schioth HB, Gloriam DE. Trends in GPCR drug discovery: new agents, targets and indications. Nat Rev Drug Discov. 2017;16(12):829–42.

2. Weis WI, Kobilka BK. The Molecular Basis of G Protein-Coupled Receptor Activation. Annu Rev Biochem. 2018;87:897–919.

3. Thompson MD, Percy ME, Cole DEC, Bichet DG, Hauser AS, Gorvin CM. G protein-coupled receptor (GPCR) gene variants and human genetic disease. Crit Rev Clin Lab Sci. 2024;61(5):317–46.

4. Eiger DS, Hicks C, Gardner J, Pham U, Rajagopal S. Location bias: A “Hidden Variable” in GPCR pharmacology. Bioessays. 2023;45(11):e2300123.

5. Kenakin T. Biased Receptor Signaling in Drug Discovery. Pharmacol Rev. 2019;71(2):267–315.

6. Gross JD, Kim DW, Zhou Y, Jansen D, Slosky LM, Clark NB, et al. Discovery of a functionally selective ghrelin receptor (GHSR(1a)) ligand for modulating brain dopamine. Proc Natl Acad Sci U S A. 2022;119(10):e2112397119.

7. Rajagopal S, Shenoy SK. GPCR desensitization: Acute and prolonged phases. Cell Signal. 2018;41:9–16.

8. Yanagi S, Sato T, Kangawa K, Nakazato M. The Homeostatic Force of Ghrelin. Cell Metab. 2018;27(4):786–804.

9. Gross JD, Zhou Y, Barak LS, Caron MG. Ghrelin receptor signaling in health and disease: a biased view. Trends Endocrinol Metab. 2023;34(2):106–18.

10. Evron T, Peterson SM, Urs NM, Bai Y, Rochelle LK, Caron MG, et al. G Protein and beta-arrestin signaling bias at the ghrelin receptor. J Biol Chem. 2014;289(48):33442–55.

11. Hedegaard MA, Holst B. The Complex Signaling Pathways of the Ghrelin Receptor. Endocrinology. 2020;161(4).

12. Mende F, Hundahl C, Plouffe B, Skov LJ, Sivertsen B, Madsen AN, et al. Translating biased signaling in the ghrelin receptor system into differential in vivo functions. Proc Natl Acad Sci U S A. 2018;115(43):E10255–E64.

13. Nagi K, Habib AM. Biased signaling: A viable strategy to drug ghrelin receptors for the treatment of obesity. Cell Signal. 2021;83:109976.

14. Toth K, Nagi K, Slosky LM, Rochelle L, Ray C, Kaur S, et al. Encoding the beta-Arrestin Trafficking Fate of Ghrelin Receptor GHSR1a: C-Tail-Independent Molecular Determinants in GPCRs. ACS Pharmacol Transl Sci. 2019;2(4):230–46.

15. Liu H, Sun D, Myasnikov A, Damian M, Baneres JL, Sun J, et al. Structural basis of human ghrelin receptor signaling by ghrelin and the synthetic agonist ibutamoren. Nat Commun. 2021;12(1):6410.

16. Qin J, Cai Y, Xu Z, Ming Q, Ji SY, Wu C, et al. Molecular mechanism of agonism and inverse agonism in ghrelin receptor. Nat Commun. 2022;13(1):300.

17. Shiimura Y, Horita S, Hamamoto A, Asada H, Hirata K, Tanaka M, et al. Structure of an antagonist-bound ghrelin receptor reveals possible ghrelin recognition mode. Nat Commun. 2020;11(1):4160.

18. Wang Y, Guo S, Zhuang Y, Yun Y, Xu P, He X, et al. Molecular recognition of an acyl-peptide hormone and activation of ghrelin receptor. Nat Commun. 2021;12(1):5064.

19. Gimpl G. Interaction of G protein coupled receptors and cholesterol. Chem Phys Lipids. 2016;199:61–73.

20. Casiraghi M, Damian M, Lescop E, Point E, Moncoq K, Morellet N, et al. Functional Modulation of a G Protein-Coupled Receptor Conformational Landscape in a Lipid Bilayer. J Am Chem Soc. 2016;138(35):11170–5.

21. Simons K, Toomre D. Lipid rafts and signal transduction. Nat Rev Mol Cell Biol. 2000;1(1):31–9.

22. Moreau CJ, Audic G, Lemel L, Garcia-Fernandez MD, Niescierowicz K. Interactions of cholesterol molecules with GPCRs in different states: A comparative analysis of GPCRs’ structures. Biochim Biophys Acta Biomembr. 2023;1865(3):184100.

23. Kim DM, Lee JH, Pan Q, Han HW, Shen Z, Eshghjoo S, et al. Nutrient-sensing growth hormone secretagogue receptor in macrophage programming and meta-inflammation. Mol Metab. 2024;79:101852.

24. Drube J, Haider RS, Matthees ESF, Reichel M, Zeiner J, Fritzwanker S, et al. GPCR kinase knockout cells reveal the impact of individual GRKs on arrestin binding and GPCR regulation. Nat Commun. 2022;13(1):540.

25. Matthees ESF, Filor JC, Jaiswal N, Reichel M, Youssef N, D’Uonnolo G, et al. GRK specificity and Gbetagamma dependency determines the potential of a GPCR for arrestin-biased agonism. Commun Biol. 2024;7(1):802.

26. Kawakami K, Yanagawa M, Hiratsuka S, Yoshida M, Ono Y, Hiroshima M, et al. Heterotrimeric Gq proteins act as a switch for GRK5/6 selectivity underlying beta-arrestin transducer bias. Nat Commun. 2022;13(1):487.

27. Sarma P, Carino CMC, Seetharama D, Pandey S, Dwivedi-Agnihotri H, Rui X, et al. Molecular insights into intrinsic transducer-coupling bias in the CXCR4-CXCR7 system. Nat Commun. 2023;14(1):4808.

28. Pack TF, Orlen MI, Ray C, Peterson SM, Caron MG. The dopamine D2 receptor can directly recruit and activate GRK2 without G protein activation. J Biol Chem. 2018;293(16):6161–71.

29. Zhao LH, Lin J, Ji SY, Zhou XE, Mao C, Shen DD, et al. Structure insights into selective coupling of G protein subtypes by a class B G protein-coupled receptor. Nat Commun. 2022;13(1):6670.

30. Marion S, Oakley RH, Kim KM, Caron MG, Barak LS. A beta-arrestin binding determinant common to the second intracellular loops of rhodopsin family G protein-coupled receptors. J Biol Chem. 2006;281(5):2932–8.

31. Toth AD, Szalai B, Kovacs OT, Garger D, Prokop S, Soltesz-Katona E, et al. G protein-coupled receptor endocytosis generates spatiotemporal bias in beta-arrestin signaling. Sci Signal. 2024;17(842):eadi0934.

32. Olsen RHJ, DiBerto JF, English JG, Glaudin AM, Krumm BE, Slocum ST, et al. TRUPATH, an open-source biosensor platform for interrogating the GPCR transducerome. Nat Chem Biol. 2020;16(8):841–9.

33. Damian M, Marie J, Leyris JP, Fehrentz JA, Verdie P, Martinez J, et al. High constitutive activity is an intrinsic feature of ghrelin receptor protein: a study with a functional monomeric GHS-R1a receptor reconstituted in lipid discs. J Biol Chem. 2012;287(6):3630–41.

34. Wan Q, Okashah N, Inoue A, Nehme R, Carpenter B, Tate CG, et al. Mini G protein probes for active G protein-coupled receptors (GPCRs) in live cells. J Biol Chem. 2018;293(19):7466–73.

35. Du Y, Duc NM, Rasmussen SGF, Hilger D, Kubiak X, Wang L, et al. Assembly of a GPCR-G Protein Complex. Cell. 2019;177(5):1232–42 e11.

36. Franke RR, Konig B, Sakmar TP, Khorana HG, Hofmann KP. Rhodopsin mutants that bind but fail to activate transducin. Science. 1990;250(4977):123–5.

37. Rouault AAJ, Buscaglia P, Sebag JA. MRAP2 inhibits beta-arrestin recruitment to the ghrelin receptor by preventing GHSR1a phosphorylation. J Biol Chem. 2022;298(6):102057.

38. Koch WJ, Inglese J, Stone WC, Lefkowitz RJ. The binding site for the beta gamma subunits of heterotrimeric G proteins on the beta-adrenergic receptor kinase. J Biol Chem. 1993;268(11):8256–60.

39. Moss J, Vaughan M. ADP-ribosylation of guanyl nucleotide-binding regulatory proteins by bacterial toxins. Adv Enzymol Relat Areas Mol Biol. 1988;61:303–79.

40. Broglio F, Benso A, Gottero C, Prodam F, Grottoli S, Tassone F, et al. Effects of glucose, free fatty acids or arginine load on the GH-releasing activity of ghrelin in humans. Clin Endocrinol (Oxf). 2002;57(2):265–71.

41. Wingler LM, McMahon C, Staus DP, Lefkowitz RJ, Kruse AC. Distinctive Activation Mechanism for Angiotensin Receptor Revealed by a Synthetic Nanobody. Cell. 2019;176(3):479–90 e12.

42. Liu X, Masoudi A, Kahsai AW, Huang LY, Pani B, Staus DP, et al. Mechanism of beta(2)AR regulation by an intracellular positive allosteric modulator. Science. 2019;364(6447):1283–7.

43. Powers AS, Khan A, Paggi JM, Latorraca NR, Souza S, Salvo JD, et al. A non-canonical mechanism of GPCR activation. bioRxiv. 2023.

44. Casiraghi M, Wang H, Brennan PC, Habrian C, Hubner H, Schmidt MF, et al. Structure and dynamics determine G protein coupling specificity at a class A GPCR. Sci Adv. 2025;11(12):eadq3971.

45. Oh P, Schnitzer JE. Segregation of heterotrimeric G proteins in cell surface microdomains. G(q) binds caveolin to concentrate in caveolae, whereas G(i) and G(s) target lipid rafts by default. Mol Biol Cell. 2001;12(3):685–98.

46. Waheed AA, Jones TL. Hsp90 interactions and acylation target the G protein Galpha 12 but not Galpha 13 to lipid rafts. J Biol Chem. 2002;277(36):32409–12.

47. Zheng H, Chu J, Qiu Y, Loh HH, Law PY. Agonist-selective signaling is determined by the receptor location within the membrane domains. Proc Natl Acad Sci U S A. 2008;105(27):9421–6.

48. Slosky LM, Caron MG, Barak LS. Biased Allosteric Modulators: New Frontiers in GPCR Drug Discovery. Trends Pharmacol Sci. 2021;42(4):283–99.

49. Barak LS, Oakley RH, Laporte SA, Caron MG. Constitutive arrestin-mediated desensitization of a human vasopressin receptor mutant associated with nephrogenic diabetes insipidus. Proc Natl Acad Sci U S A. 2001;98(1):93–8.

50. Zheng K, Smith JS, Eiger DS, Warman A, Choi I, Honeycutt CC, et al. Biased agonists of the chemokine receptor CXCR3 differentially signal through Galpha(i):beta-arrestin complexes. Sci Signal. 2022;15(726):eabg5203.

51. Snyder JC, Rochelle LK, Ray C, Pack TF, Bock CB, Lubkov V, et al. Inhibiting clathrin-mediated endocytosis of the leucine-rich G protein-coupled receptor-5 diminishes cell fitness. J Biol Chem. 2017;292(17):7208–22.

52. Smith JS, Alagesan P, Desai NK, Pack TF, Wu JH, Inoue A, et al. C-X-C Motif Chemokine Receptor 3 Splice Variants Differentially Activate Beta-Arrestins to Regulate Downstream Signaling Pathways. Mol Pharmacol. 2017;92(2):136–50.

