## Supplementary Data for "Rare variant in intracellular loop-2 of the ghrelin receptor reveals novel mechanisms of GPCR biased signaling and trafficking"

| ***Figure/Panel*** | ***Description & Units*** | ***Condition/***  ***Treatment*** | ***Mean ± SEM*** | ***t/F-Statistic &***  ***P-value*** | ***Multiple Comparisons*** |
| --- | --- | --- | --- | --- | --- |
| 1b | On-Cell ELISA  (% WT-pcDNA-vehicle) | WT-background  L149P-background  WT+Dyn^K44A^-background  L149P+Dyn^K44A^-background  WT+pcDNA-vehicle  L149P+pcDNA-vehicle  WT+Dyn^K44A^-vehicle  L149P+Dyn^K44A^-vehicle  WT+pcDNA-hGhrelin  L149P+pcDNA-hGhrelin  WT+Dyn^K44A^-hGhrelin  L149P+Dyn^K44A^-hGhrelin  WT+pcDNA-MK0677  L149P+pcDNA-MK0677  WT+Dyn^K44A^-MK0677  L149P+Dyn^K44A^-MK0677 | 0.0001±2.844  3.351±3.696  5.478±2.887  5.174±3.493  100.00±2.224  234.35±3.423  250.09±11.149  363.04±17.313  69.411±3.099  168.20±4.832  245.68±10.325  348.57±17.054  66.004±4.229  167.38±10.747  252.53±9.713  345.38±15.487 | *Interaction*:  F(9,215)=41.34  P<0.0001****  *Condition*:  F(3,215)=381.5  P<0.0001****  *Treatment*:  F(3,215)=498.0  P<0.0001**** | *Holm-Sidak Post-Hoc Test (selected comparisons)*  WT+pcDNA-vehicle vs L149P+pcDNA-vehicle  P<0.0001****  WT+pcDNA-vehicle vs WT+Dyn^K44A^-vehicle  P<0.0001****  WT+pcDNA-vehicle vs WT+pcDNA-hGhrelin  P=0.068  WT+pcDNA-vehicle vs WT+pcDNA-MK0677  P=0.0356*  L149P+pcDNA-vehicle vs WT+Dyn^K44A^-vehicle  P=0.697  L149P+pcDNA-vehicle vs L149P+pcDNA-hGhrelin  P<0.0001****  L149P+pcDNA-vehicle vs L149P+pcDNA-MK0677  P<0.0001****  WT+Dyn^K44A^-vehicle vs WT+Dyn^K44A^-hGhrelin  P=0.944  WT+Dyn^K44A^-vehicle vs WT+Dyn^K44A^-MK0677  P=0.944  L149P+Dyn^K44A^-vehicle vs L149P+Dyn^K44A^-hGhrelin  P=0.697  L149P+Dyn^K44A^-vehicle vs L149P+Dyn^K44A^-MK0677  P=0.601  L149P+pcDNA-vehicle vs L149P+Dyn^K44A^-vehicle  P<0.0001**** |
| 1c | In-Cell ELISA  (% WT control) | WT^100ng^  L149P^50ng^  L149P^100ng^ | 100.0±3.764  110.0±3.398  241.2±7.852 | F(2,44)=235.6  P<0.0001**** | *Tukey’s Post-Hoc Test*  WT^100ng^ vs L149P^50ng^  P=0.268  WT^100ng^ vs L149P^100ng^  P<0.0001****  L149P^50ng^ vs L149P^100ng^  P<0.0001**** |
| 1e | Basal CAAX & MyrPalm  (Δ net BRET; relative to empty vector control) | CAAX-WT  CAAX-L149P  MyrPalm-WT  MyrPalm-L149P | 0.717±0.013  0.902±0.021  1.187±0.020  1.327±0.034 | *Interaction*:  F(1,144)=1.029  P=0.312  *bBRET Sensor*:  F(1,144)=411.2  P<0.0001****  *Receptor*:  F(1,144)=54.12  P<0.0001**** | *Tukey’s Post-Hoc Test*  CAAX-WT vs CAAX-L149P  P<0.0001****  CAAX-WT vs MyrPalm-WT  P<0.0001****  CAAX-WT vs MyrPalm-L149P  P<0.0001****  CAAX-L149P vs MyrPalm-WT  P<0.0001****  CAAX-L149P vs MyrPalm-L149P  P<0.0001****  MyrPalm-WT vs MyrPalm-L149P  0.0005*** |
| 1f | Basal Lipid Raft (MyrPalm) Enrichment  (ΔΔ net BRET; ratio of MyrPalm [Δ net BRET] to CAAX [Δ net BRET]) | WT  L149P | 1.647±0.040  1.419±0.044 | t(22)=3.803  0.0005*** | N/A |
| 1g | Lipid Raft (MyrPalm) Enrichment after MβCD  (ΔΔ net BRET; ratio of MyrPalm [Δ net BRET] to CAAX [Δ net BRET]) | WT (post-MβCD)  L149P (post-MβCD) | 1.891±0.036  2.091±0.084 | t(30)=2.183  P=0.037* | N/A |
| 1h | Basal 2xFYVE  (Δ net BRET; relative to empty vector control) | WT  L149P | 0.288±0.011  0.215±0.008 | t(70)=5.194  P<0.0001**** | N/A |
| 1i | Basal PM:Endosome Ratio (CAAX/MyrPalm:2xFYVE)  (ΔΔ net BRET; ratio of CAAX:2xFYVE vs ratio of MyrPalm:2xFYVE) | CAAX:2xFYVE-WT  CAAX:2xFYVE-L149P  MyrPalm:2xFYVE-WT  MyrPalm:2xFYVE-L149P | 2.480±0.059  4.324±0.103  4.121±0.081  6.196±0.147 | *Interaction*:  F(1,88)=1.296  P=0.258  *Ratio*:  F(1,88)=299.0  P<0.0001****  *Receptor*:  F(1,88)=372.4  P<0.0001**** | *Tukey’s Post-Hoc Test*  CAAX:2xFYVE-WT vs CAAX:2xFYVE-L149P  P<0.0001****  CAAX:2xFYVE-WT vs MyrPalm:2xFYVE-WT  P<0.0001****  CAAX:2xFYVE-WT vs MyrPalm:2xFYVE-L149P  P<0.0001****  CAAX:2xFYVE-L149P vs MyrPalm:2xFYVE-WT  P=0.491  CAAX:2xFYVE-L149P vs MyrPalm:2xFYVE-L149P  P<0.0001****  MyrPalm:2xFYVE-WT vs MyrPalm:2xFYVE-L149P  P<0.0001**** |
| 1j | PM:Endosome Ratio (CAAX/MyrPalm:2xFYVE) after MβCD  (ΔΔ net BRET; ratio of CAAX:2xFYVE vs ratio of MyrPalm:2xFYVE) | CAAX:2xFYVE-WT (post-MβCD)  CAAX:2xFYVE-L149P (post-MβCD)  MyrPalm:2xFYVE-WT (post-MβCD)  MyrPalm:2xFYVE-L149P (post-MβCD) | 17.620±0.522  7.480±1.110  33.311±0.639  15.639±0.631 | *Interaction*:  F(1,60)=24.55  P<0.0001****  *Ratio*:  F(1,60)=245.2  P<0.0001****  *Receptor*:  F(1,60)=334.8  P<0.0001**** | *Tukey’s Post-Hoc Test*  CAAX:2xFYVE-WT vs CAAX:2xFYVE-L149P  P<0.0001****  CAAX:2xFYVE-WT vs MyrPalm:2xFYVE-WT  P<0.0001****  CAAX:2xFYVE-WT vs MyrPalm:2xFYVE-L149P  P<0.0001****  CAAX:2xFYVE-L149P vs MyrPalm:2xFYVE-WT  P=0.263  CAAX:2xFYVE-L149P vs MyrPalm:2xFYVE-L149P  P<0.0001****  MyrPalm:2xFYVE-WT vs MyrPalm:2xFYVE-L149P  P<0.0001**** |
| 1k | CAAX (non-raft): hGhrelin Internalization  (area under the curve [AUC]; negative area peaks) | CAAX-WT-hGhrelin  CAAX-L149P-hGhrelin | 0.904±0.675  1.036±0.531 | t(598)=0.152  P=0.878 | N/A |
| 1l | MyrPalm (lipid raft): hGhrelin Internalization  (area under the curve [AUC]; negative area peaks) | MyrPalm-WT-hGhrelin  MyrPalm-L149P-hGhrelin | 20.820±0.970  7.372±0.802 | t(598)=10.68  P<0.0001**** | N/A |
| 1m | 2xFYVE: hGhrelin Endosomal Translocation  (area under the curve [AUC]; positive area peaks) | 2xFYVE-WT-hGhrelin  2xFYVE-L149P-hGhrelin | 1.374±0.250  7.055±0.334 | t(598)=13.59  P<0.0001**** | N/A |
| 1n | MyrPalm (lipid raft):  hGhrelin Internalization after MβCD | MyrPalm-WT-hGhrelin (vehicle)  MyrPalm-L149P-hGhrelin (vehicle)  MyrPalm-WT-hGhrelin (MβCD)  MyrPalm-L149P-hGhrelin (MβCD) | -0.312±0.013  -0.139±0.014  -0.156±0.010  -0.103±0.012 | *Interaction*:  F(1,60)=23.69  P<0.0001****  *MβCD*:  F(1,60)=61.03  P<0.0001****  *Receptor*:  F(1,60)=83.46  P<0.0001**** | *Tukey’s Post-Hoc Test*  WT-vehicle vs L149P-vehicle  P<0.0001****  WT-vehicle vs WT-MβCD  P<0.0001****  WT-vehicle vs L149P-MβCD  P<0.0001****  L149P-vehicle vs WT-MβCD  P=0.785  L149P-vehicle vs L149P-MβCD  P=0.170  WT-MβCD vs L149P-MβCD  P=0.019* |
| 1o | 2xFYVE:  hGhrelin Endosomal Translocation after MβCD | 2xFYVE-WT-hGhrelin (vehicle)  2xFYVE-L149P-hGhrelin (vehicle)  2xFYVE-WT-hGhrelin (MβCD)  2xFYVE-L149P-hGhrelin (MβCD) | 0.040±0.003  0.107±0.006  0.042±0.005  0.084±0.005 | *Interaction*:  F(1,60)=6.261  P=0.0151*  *MβCD*:  F(1,60)=4.499  P=0.038*  *Receptor*:  F(1,60)=122.7  P<0.0001**** | *Tukey’s Post-Hoc Test*  WT-vehicle vs L149P-vehicle  P<0.0001****  WT-vehicle vs WT-MβCD  P=0.993  WT-vehicle vs L149P-MβCD  P<0.0001****  L149P-vehicle vs WT-MβCD  P<0.0001****  L149P-vehicle vs L149P-MβCD  P=0.009**  WT-MβCD vs L149P-MβCD  P<0.0001**** |

N/A: Not Applicable

**Supplementary Table 1.** Data and statistics from main **Figure 1**.

| ***Figure/Panel*** | ***Description & Units*** | ***Condition/***  ***Treatment*** | ***Mean ± SEM*** | ***t/F-Statistic &***  ***P-value*** | ***Multiple Comparisons*** | ***pEC_50_/K_m_ ± SEM*** | ***F-Statistic &***  ***P-value*** |
| --- | --- | --- | --- | --- | --- | --- | --- |
| 2b | hβarr1:  Basal Activity  (net BRET) | WT  L149P | 0.884±0.002  0.875±0.004 | t(34)=1.591 p=0.120 | N/A | N/A  N/A | N/A |
| 2c | hβarr1:  hGhrelin C/R  (% WT) | WT  L149P | N/A  N/A | N/A | N/A | 8.212±0.051  8.108±0.097 | F(1,100)=0.766  P=0.383 |
| 2d | hβarr1:  hGhrelin Kinetics  (net BRET) | WT  L149P | N/A  N/A | N/A | N/A | 15.136±1.486  20.323±4.066 | F(1,227)=1.714  P=0.191 |
| 2e | hβarr1:  MK0677 C/R  (% WT) | WT  L149P | N/A  N/A | N/A | N/A | 8.210±0.044  8.118±0.053 | F(1,94)=1.162  P=0.283 |
| 2f | hβarr1:  MK0677 Kinetics  (net BRET) | WT  L149P | N/A  N/A | N/A | N/A | 7.655±1.090  5.738±0.803 | F(1,216)=0.988  P=0.321 |
| 2g | hβarr1:  Agonist E_max_  (% WT) | WT-hGhrelin  L149P-hGhrelin  WT-MK0677  L149P-MK0677 | 100±2.221  47.867±2.272  100±1.855  43.428±1.194 | *Interaction*:  F(1,212)=1.286  P-0.2561  *Agonist*:  F(1,212)=0.247  P=0.247  *Receptor*:  F(1,212)=790.5  P<0.0001**** | *Tukey’s Post-Hoc*:  WT-hGhrelin vs L149P-hGhrelin  P<0.0001****  WT-hGhrelin vs WT-MK0677  P<0.999  WT-hGhrelin vs L149P-MK0677  P<0.0001****  L149P-hGhrelin vs WT-MK0677  P<0.0001****  L149P-hGhrelin vs L149P-MK0677  P=0.368  WT-MK0677 vs L149P-MK0677  P<0.0001**** | N/A  N/A  N/A  N/A | N/A |
| 2i | hβarr2:  Basal Activity  (net BRET) | WT  L149P | 1.309±0.008  1.103±0.008 | T(34)=18.03  P<0.0001**** | N/A | N/A  N/A | N/A |
| 2j | hβarr2:  hGhrelin C/R  (% WT) | WT  L149P | N/A  N/A | N/A | N/A | 7.966±0.079  7.947±0.058 | F(1,100)=0.033  P=0.854 |
| 2k | hβarr2:  hGhrelin Kinetics  (net BRET) | WT  L149P | N/A  N/A | N/A | N/A | 7.189±1.001  9.011±1.382 | F(1,230)=1.108  P=0.293 |
| 2l | hβarr2:  MK0677 C/R  (% WT) | WT  L149P | N/A  N/A | N/A | N/A | 8.311±0.030  8.190±0.039 | F(1,100)=5.977  P=0.016* |
| 2m | hβarr2:  MK0677 Kinetics  (net BRET) | WT  L149P | N/A  N/A | N/A | N/A | 2.481±0.487  3.260±0.381 | F(1,230)=1.363  P=0.2442 |
| 2n | hβarr2:  Agonist E_max_  (% WT) | WT-hGhrelin  L149P-hGhrelin  WT-MK0677  L149P-MK0677 | 100±4.363  66.495±2.254  100±1.132  76.127±1.144 | *Interaction*:  F(1,212)=3.405  P=0.066  *Agonist*:  F(1,212)=3.348  P=0.068  *Receptor*:  F(1,212)=120.0  P<0.0001**** | *Tukey’s Post-Hoc*:  WT-hGhrelin vs L149P-hGhrelin  P<0.0001****  WT-hGhrelin vs WT-MK0677  P>0.999  WT-hGhrelin vs L149P-MK0677  P<0.0001****  L149P-hGhrelin vs WT-MK0677  P<0.0001****  L149P-hGhrelin vs L149P-MK0677  P=0.048*  WT-MK0677 vs L149P-MK0677  P<0.0001**** | N/A  N/A  N/A  N/A | N/A |
| 2p | hβarr1±Dyn^K44A^:  Basal Activity  (net BRET) | WT-pcDNA  L149P-pcDNA  WT-Dyn^K44A^  L149P-Dyn^K44A^ | 0.844±0.006  0.849±0.005  0.837±0.002  0.839±0.003 | *Interaction*:  F(1,40)=0.259  P=0.613  *Dyn^K44A^*:  F(1,40)=3.773  P=0.0592  *Receptor*:  F(1,40)=0.575  P=0.452 | N/A | N/A  N/A  N/A  N/A | N/A |
| 2q | hβarr1±Dyn^K44A^:  hGhrelin C/R  (% WT-pcDNA) | WT-pcDNA  L149P-pcDNA  WT-Dyn^K44A^  L149P-Dyn^K44A^ | N/A  N/A  N/A  N/A | N/A | N/A | 7.800±0.177  7.691.0.423  7.629±0.140  7.501±0.359 | F(3,320)=0.192  P-0.901 |
| 2r | hβarr1±Dyn^K44A^:  hGhrelin E_max_  (% WT-pcDNA) | WT-pcDNA  L149P-pcDNA  WT-Dyn^K44A^  L149P-Dyn^K44A^ | 100±3.149  36.573±2.300  34.895±2.300  7.636±2.064 | *Interaction*:  F(1,388)=41.45  P<0.0001****  *Dyn^K44A^*:  F(1,388)=280.2  P<0.0001****  *Receptor*:  F(1,388)=260.5  P<0.0001**** | *Tukey’s Post-Hoc*:  WT-pcDNA vs L149P-pcDNA  P<0.0001****  WT-pcDNA vs WT-Dyn^K44A^  P<0.0001****  WT-pcDNA vs L149P-Dyn^K44A^  P<0.0001****  L149P-pcDNA vs WT-Dyn^K44A^  P-0.974  L149P-pcDNA vs L149P-Dyn^K44A^  P<0.0001****  WT-Dyn^K44A^ vs L149-Dyn^K44A^  P<0.0001**** | N/A  N/A  N/A  N/A | N/A |
| 2s | hβarr2±Dyn^K44A^:  Basal Activity  (net BRET) | WT-pcDNA  L149P-pcDNA  WT-Dyn^K44A^  L149P-Dyn^K44A^ | 1.261±0.014  1.074±0.007  1.062±0.014  1.050±0.012 | *Interaction*:  F(1,52)=54.02  P<0.0001****  *Dyn^K44A^*:  F(1,52)=88.73  P<0.0001****  *Receptor*:  F(1,52)=70.00  P<0.0001**** | *Tukey’s Post-Hoc*:  WT-pcDNA vs L149P-pcDNA  P<0.0001****  WT-pcDNA vs WT-Dyn^K44A^  P<0.0001****  WT-pcDNA vs L149P-Dyn^K44A^  P<0.0001****  L149P-pcDNA vs WT-Dyn^K44A^  P=0.878  L149P-pcDNA vs L149P-Dyn^K44A^  P=0.466  WT-Dyn^K44A^ vs L149-Dyn^K44A^  P=0.889 | N/A  N/A  N/A  N/A | N/A |
| 2t | hβarr2±Dyn^K44A^:  hGhrelin C/R  (% WT-pcDNA) | WT-pcDNA  L149P-pcDNA  WT-Dyn^K44A^  L149P-Dyn^K44A^ | N/A  N/A  N/A  N/A | N/A | N/A | 8.059±0.065  8.027±0.060  8.103±0.096  7.962±0.197 | F(3,320)=0.230  P-0.874 |
| 2u | hβarr2±Dyn^K44A^:  hGhrelin E_max_  (% WT-pcDNA) | WT-pcDNA  L149P-pcDNA  WT-Dyn^K44A^  L149P-Dyn^K44A^ | 100±3.287  60.944±2.106  57.481±2.758  39.472±4.323 | *Interaction*:  F(1,388)=10.66  P-0.001**  *Dyn^K44A^*:  F(1,388)=98.55  P<0.0001****  *Receptor*:  F(1,388)=78.37  P<0.0001**** | *Tukey’s Post-Hoc*:  WT-pcDNA vs L149P-pcDNA  P<0.0001****  WT-pcDNA vs WT-Dyn^K44A^  P<0.0001****  WT-pcDNA vs L149P-Dyn^K44A^  P<0.0001****  L149P-pcDNA vs WT-Dyn^K44A^  P=0.872  L149P-pcDNA vs L149P-Dyn^K44A^  P<0.0001****  WT-Dyn^K44A^ vs L149-Dyn^K44A^  P=0.0005*** | N/A  N/A  N/A  N/A | N/A |
| 2w | hβarr1±MβCD:  Basal Activity  (net BRET) | WT-vehicle  L149P-vehicle  WT-MβCD  L149P-MβCD | 0.867±0.007  0.846±0.004  0.854±0.007  0.843±0.004 | *Interaction*:  F(1,44)=0.727  P=0.398  *MβCD*:  F(1,44)=1.914  P=0.173  *Receptor*:  F(1,44)=7.777  P=0.0078** | *Tukey’s Post-Hoc*:  WT-vehicle vs L149P-vehicle  P=0.062  WT-vehicle vs WT-MβCD  P=0.399  WT-vehicle vs L149P-MβCD  P=0.025*  L149P-vehicle vs WT-MβCD  P=0.753  L149P-vehicle vs L149P-MβCD  P=0.981  WT-MβCD vs L149-MβCD  P=0.523 | N/A  N/A  N/A  N/A | N/A |
| 2x | hβarr1± MβCD:  hGhrelin C/R  (% WT-pcDNA) | WT-vehicle  L149P-vehicle  WT-MβCD  L149P-MβCD | N/A  N/A  N/A  N/A | N/A | N/A | 7.697±0.104  7.562±0.113  7.434±0.087  7.141±0.095 | F(3,272)=4.099  P=0.0072** |
| 2y | hβarr1± MβCD:  hGhrelin E_max_  (% WT-pcDNA) | WT-vehicle  L149P-vehicle  WT-MβCD  L149P-MβCD | 100±3.602  45.516±2.895  88.964±3.738  31.883±2.615 | *Interaction*:  F(1,284)=0.160  P=0.689  *MβCD*:  F(1,284)=14.43  P=0.0002***  *Receptor*:  F(1,284)=295.1  P<0.0001*** | *Tukey’s Post-Hoc*:  WT-vehicle vs L149P-vehicle  P<0.0001****  WT-vehicle vs WT-MβCD  P=0.078  WT-vehicle vs L149P-MβCD  P<0.0001****  L149P-vehicle vs WT-MβCD  P<0.0001****  L149P-vehicle vs L149P-MβCD  P=0.017*  WT-MβCD vs L149-MβCD  P<0.0001**** | N/A  N/A  N/A  N/A | N/A |
| 2z | hβarr2±MβCD:  Basal Activity  (net BRET) | WT-vehicle  L149P-vehicle  WT-MβCD  L149P-MβCD | 1.194±0.017  1.021±0.009  1.107±0.012  0.995±0.008 | *Interaction*:  F(1,44)=6.424  P=0.014*  *MβCD*:  F(1,44)=21.33  P<0.0001****  *Receptor*:  F(1,44)=136.0  P<0.0001**** | *Tukey’s Post-Hoc*:  WT-vehicle vs L149P-vehicle  P<0.0001****  WT-vehicle vs WT-MβCD  P<0.0001****  WT-vehicle vs L149P-MβCD  P<0.0001****  L149P-vehicle vs WT-MβCD  P<0.0001****  L149P-vehicle vs L149P-MβCD  P=0.461  WT-MβCD vs L149-MβCD  P<0.0001**** | N/A  N/A  N/A  N/A | N/A |
| 2aa | hβarr2± MβCD:  hGhrelin C/R  (% WT-pcDNA) | WT-vehicle  L149P-vehicle  WT-MβCD  L149P-MβCD | N/A  N/A  N/A  N/A | N/A | N/A | 7.610±0.068  7.676±0.077  7.344±0.099  7.257±0.082 | F(3,272)=5.377  P=0.0013** |
| 2bb | hβarr2± MβCD:  hGhrelin E_max_  (% WT-pcDNA) | WT-vehicle  L149P-vehicle  WT-MβCD  L149P-MβCD | 100±1.655  79.761±2.883  86.507±4.090  57.342±2.514 | *Interaction*:  F(1,284)=1.996  P=0.158  *MβCD*:  F(1,284)=32.31  P<0.0001****  *Receptor*:  F(1,284)=61.15  P<0.0001**** | *Tukey’s Post-Hoc*:  WT-vehicle vs L149P-vehicle  0.0007***  WT-vehicle vs WT-MβCD  P=0.014*  WT-vehicle vs L149P-MβCD  P<0.0001****  L149P-vehicle vs WT-MβCD  P=0.432  L149P-vehicle vs L149P-MβCD  P<0.0001****  WT-MβCD vs L149-MβCD  P<0.0001**** | N/A  N/A  N/A  N/A | N/A |

N/A: Not Applicable

**Supplementary Table 2.** Data and statistics from main **Figure 2**.

| ***Figure/Panel*** | ***Description & Units*** | ***Condition/***  ***Treatment*** | ***Mean ± SEM*** | ***t/F-Statistic &***  ***P-value*** | ***Multiple Comparisons*** | ***pEC_50_ ± SEM*** | ***F-Statistic &***  ***P-value*** |
| --- | --- | --- | --- | --- | --- | --- | --- |
| 3b | Gα_q_ Dissociation: Constitutive  (% WT) | WT  L149P | 100.0±3.788  29.02±2.570 | t(34)=15.51  P<0.0001*** | N/A | N/A  N/A | N/A |
| 3c | Gα_q_ Dissociation: hGhrelin C/R  (% WT) | WT  L149P | N/A  N/A | N/A | N/A | 7.651±0.115  9.289±1.434 | F(1,156)=1.284  P=0.258 |
| 3d | Gα_q_ Dissociation: MK0677 C/R  (% WT) | WT  L149P | N/A  N/A | N/A | N/A | 8.282±0.064  7.170±1.643 | F(1,156)=0.784  P=0.377 |
|  | Gα_q_ Dissociation: Agonist E_max_  (% WT control) | WT-hGhrelin  L149P-hGhrelin  WT-MK0677  L149P-MK0677 | 100±4.816  -10.288±3.415  100±2.283  -9.186±6.765 | *Interaction*:  F(1,344)=0.013  P=0.907  *Agonist*:  F(1,344)=0.014  P=0.903  *Receptor*:  F(1,344)=561.1  P<0.0001**** | *Tukey’s Post-Hoc*:  WT-hGhrelin vs L149P-hGhrelin  P<0.0001****  WT-hGhrelin vs WT-MK0677  P>0.999  WT-hGhrelin vs L149P-MK0677  P<0.0001****  L149P-hGhrelin vs WT-MK0677  P<0.0001****  L149P-hGhrelin vs L149P-MK0677  P=0.998  WT-MK0677 vs L149P-MK0677  P<0.0001**** | N/A  N/A  N/A  N/A | N/A |
| 3f | mini-G_q_: Constitutive  (% WT) | WT  L149P | 100.0±3.566  3.007±1.853 | t(26)=24.14  P<0.0001**** | N/A | N/A  N/A | N/A |
| 3g | mini-G_q_: hGhrelin C/R  (% WT) | WT  L149P | N/A  N/A | N/A | N/A | 7.780±0.084  8.009±0.154 | F(1,62)=0.729  P=0.396 |
| 3h | mini-G_q_: MK0677  (% WT) | WT  L149P | N/A  N/A | N/A | N/A | 8.057±0.093  8.306±0.652 | F(1,62)=0.794  P=0.376 |
|  | mini-G_q_: Agonist E_max_  (% WT control) | WT-hGhrelin  L149P-hGhrelin  WT-MK0677  L149P-MK0677 | 100±3.544  26.825±1.414  100±3.614  28.075±0.652 | *Interaction*:  F(1,132)=0.057  P=0.810  *Agonist*:  F(1,132)=0.056  P=0.812  *Receptor*:  F(1,132)=767.0  P<0.0001**** | *Tukey’s Post-Hoc*:  WT-hGhrelin vs L149P-hGhrelin  P<0.0001****  WT-hGhrelin vs WT-MK0677  P>0.999  WT-hGhrelin vs L149P-MK0677  P<0.0001****  L149P-hGhrelin vs WT-MK0677  P<0.0001****  L149P-hGhrelin vs L149P-MK0677  P=0.986  WT-MK0677 vs L149P-MK0677  P<0.0001**** | N/A  N/A  N/A  N/A | N/A |
| 3l | Gα_q_ Dissociation ± MβCD: Constitutive  (% WT-vehicle) | WT-vehicle  WT-MβCD  L149P-vehicle  L149P-MβCD | 100±7.198  85.999±7.953  14.305±9.237  23.075±8.581 | *Interaction*:  F(1,44)=1.892  P=0.1759  *M*β*CD*:  F(1,44)=0.099  P=0.753  *Receptor*:  F(1,44)=80.61  P<0.0001**** | *Sidak’s Post-Hoc*  *Vehicle: WT vs L149P*  P<0.0001****  *MβCD: WT vs L149P*  P<0.0001**** | N/A  N/A | N/A |
| 3m | Gα_q_ Dissociation ± MβCD: hGhrelin C/R  (% WT-vehicle) | WT-vehicle  WT-MβCD  L149P-vehicle  L149P-MβCD | N/A  N/A | N/A | N/A | 8.226±0.056  7.811±0.075  UD  UD | F(3,272)=6.680  P=0.0002*** |
| 3n | Gα_q_ Dissociation ± MβCD: hGhrelin Emax  (% WT control) | WT-vehicle  WT-MβCD  L149P-vehicle  L149P-MβCD | 100±2.361  98.91±3.669  0.573±2.064  UD | *Interaction*:  F(1,284)=0.009  P=0.921  *M*β*CD*:  F(1,284)=0.112  P=0.737  *Receptor*:  F(1,284)=1687  P<0.0001**** | *Sidak’s Post-Hoc*  *Vehicle: WT vs L149P*  P<0.0001****  *MβCD: WT vs L149P*  P<0.0001**** | N/A  N/A | N/A |
| 3p | Gα_13_ Dissociation ± MβCD: Constitutive  (% WT-vehicle) | WT-vehicle  WT-MβCD  L149P-vehicle  L149P-MβCD | 100±19.771  183.390±27.351  87.788±11.296  125.621±18.162 | *Interaction*:  F(1,43)=1.295  P=0.261  *M*β*CD*:  F(1,43)=9.170  P=0.004**  *Receptor*:  F(1,43)=3.056  P=0.087 | *Sidak’s Post-Hoc*  *WT: vehicle vs MβCD*  P=0.011*  *L149P: vehicle vs MβCD*  P=0.333 | N/A  N/A | N/A |
| 3q | Gα_13_ Dissociation ± MβCD: hGhrelin C/R  (% WT-vehicle) | WT-vehicle  WT-MβCD  L149P-vehicle  L149P-MβCD | N/A  N/A | N/A | N/A | 6.887±0.116  6.620±0.209  UD  UD | F(3,276)=0.815  P=0.4862 |
| 3r | Gα_13_ Dissociation ± MβCD: hGhrelin Emax  (% WT control) | WT-vehicle  WT-MβCD  L149P-vehicle  L149P-MβCD | 100±6.981  75.491±9.568  10.277±3.649  5.704±2.663 | *Interaction*:  F(1,284)=2.473  P=0.116  *M*β*CD*:  F(1,284)=0.022  P=0.022*  *Receptor*:  F(1,284)=158.3  P<0.0001**** | *Tukey’s Post-Hoc*:  WT-vehicle vs L149P-vehicle  P<0.0001****  WT-vehicle vs WT-MβCD  P=0.033*  WT-vehicle vs L149P-MβCD  P<0.0001****  L149P-vehicle vs WT-MβCD  P<0.0001****  L149P-vehicle vs L149P-MβCD  P=0.956  WT-MβCD vs L149P-MβCD  P<0.0001**** | N/A  N/A | N/A |
| 3t | Gα_i2_ Dissociation ± MβCD: Constitutive  (% WT-vehicle) | WT-vehicle  WT-MβCD  L149P-vehicle  L149P-MβCD | 100±5.109  109.549±4.638  33.455±3.921  35.664±4.046 | *Interaction*:  F(1,44)=0.679  P=0.414  *M*β*CD*:  F(1,44)=1.742  P=0.193  *Receptor*:  F(1,44)=248.5  P<0.0001**** | *Sidak’s Post-Hoc*  *Vehicle: WT vs L149P*  P<0.0001****  *MβCD: WT vs L149P*  P<0.0001**** | N/A  N/A | N/A |
| 3u | Gα_i2_ Dissociation ± MβCD: hGhrelin C/R  (% WT-vehicle) | WT-vehicle  WT-MβCD  L149P-vehicle  L149P-MβCD | N/A  N/A | N/A | N/A | 7.095±0.138  6.755±0.242  7.078±0.359  6.539±0.755 | F(3,276)=0.642  P=0.588 |
| 3v | Gα_i2_ Dissociation ± MβCD: hGhrelin Emax  (% WT control) | WT-vehicle  WT-MβCD  L149P-vehicle  L149P-MβCD | 100±7.567  92.445±11.690  38.661±8.841  45.672±25.46 | *Interaction*:  F(1,284)=0.231  P=0.631  *M*β*CD*:  F(1,284)=0.0003  P=0.985  *Receptor*:  F(1,284)=12.70  P<0.0004*** | *Sidak’s Post-Hoc*  *Vehicle: WT vs L149P*  P=0.009**  *MβCD: WT vs L149P*  P=0.059 | N/A  N/A | N/A |
| 3x | Gα_oA_ Dissociation ± MβCD: Constitutive  (% WT-vehicle) | WT-vehicle  WT-MβCD  L149P-vehicle  L149P-MβCD | 100±4.006  113.088±5.657  25.573±2.627  33.351±3.305 | *Interaction*:  F(1,44)=0.399  P=0.530  *M*β*CD*:  F(1,44)=6.723  P=0.012*  *Receptor*:  F(1,44)=359.9  P<0.0001**** | *Tukey’s Post-Hoc*:  WT-vehicle vs L149P-vehicle  P<0.0001****  WT-vehicle vs WT-MβCD  P=0.118  WT-vehicle vs L149P-MβCD  P<0.0001****  L149P-vehicle vs WT-MβCD  P<0.0001****  L149P-vehicle vs L149P-MβCD  P=0.514  WT-MβCD vs L149P-MβCD  P<0.0001**** | N/A  N/A | N/A |
| 3y | Gα_oA_ Dissociation ± MβCD: hGhrelin C/R  (% WT-vehicle) | WT-vehicle  WT-MβCD  L149P-vehicle  L149P-MβCD | N/A  N/A | N/A | N/A | 7.374±0.077  6.972±0.103  7.108±0.174  7.236±0.174 | F(3,270)=2.870  P=0.036* |
| 3z | Gα_oA_ Dissociation ± MβCD: hGhrelin Emax  (% WT control) | WT-vehicle  WT-MβCD  L149P-vehicle  L149P-MβCD | 100±3.593  78.040±4.640  59.195±5.390  53.051±4.650 | *Interaction*:  F(1,284)=2.939  P=0.087  *M*β*CD*:  F(1,284)=9.279  P=0.002**  *Receptor*:  F(1,284)=50.85  P<0.0001**** | *Tukey’s Post-Hoc*:  WT-vehicle vs L149P-vehicle  P<0.0001****  WT-vehicle vs WT-MβCD  P=0.004**  WT-vehicle vs L149P-MβCD  P<0.0001****  L149P-vehicle vs WT-MβCD  P<0.021*  L149P-vehicle vs L149P-MβCD  P=0.782  WT-MβCD vs L149P-MβCD  P<0.0009*** | N/A  N/A | N/A |

N/A: Not Applicable; UD: Undeterminable

**Supplementary Table 4.** Data and statistics from main **Figure 3**.

| ***Figure/Panel*** | ***Description & Units*** | ***Condition/***  ***Treatment*** | ***Mean ± SEM*** | ***t/F-Statistic &***  ***P-value*** | ***Multiple Comparisons*** | ***pEC_50_ ± SEM*** | ***F-Statistic &***  ***P-value*** |
| --- | --- | --- | --- | --- | --- | --- | --- |
| 5b | WT-hβarr1: Constitutive  (Δ net BRET; % WT-vehicle) | vehicle  Go6983  Cmp101  Go6983+Cmp101 | 100±3.399  101.3±8.922  123.8±7.541  73.44±7.129 | F(3,68)=5.572  P=0.0018** | *Tukey’s Post Hoc Test*  Vehicle vs Go6983  P=0.999  Vehicle vs Cmp101  P=0.186  Vehicle vs Go6983+Cmp101  P=0.115  Go6983 vs Cmp101  0.229  Go6983 vs Go6983+Cmp101  0.090  Cmp101 vs Go6983+Cmp101  P=0.0007*** | N/A  N/A  N/A  N/A | N/A |
| 5c | WT-hβarr1: hGhrelin C/R  (Δ net BRET; % WT-vehicle) | vehicle  Go6983  Cmp101  Go6983+Cmp101 | N/A  N/A  N/A  N/A | N/A | N/A | 8.006±0.041  7.947±0.083  7.881±0.239  UD | F(3,332)=0.513  P=0.673 |
| 5d | WT-hβarr1: hGhrelin E_max_  (E_max_; % WT-vehicle) | vehicle  Go6983  Cmp101  Go6983+Cmp101 | 100±2.732  71.415±3.352  17.220±2.307  UD | F(3,344)=299.0  P<0.0001**** | *Tukey’s Post Hoc Test*  Vehicle vs Go6983  P<0.0001****  Vehicle vs Cmp101  P<0.0001****  Vehicle vs Go6983+Cmp101  P<0.0001****  Go6983 vs Cmp101  P<0.0001****  Go6983 vs Go6983+Cmp101  P<0.0001****  Cmp101 vs Go6983+Cmp101  P=0.0008*** | N/A  N/A  N/A  N/A | N/A |
| 5e | L149P-hβarr1: Constitutive  (Δ net BRET; % WT-vehicle) | vehicle  Go6983  Cmp101  Go6983+Cmp101 | 82.43±5.979  95.68±8.421  103.2±9.163  75.58±7.059 | F(3,68)=2.453  P=0.070* | *Tukey’s Post Hoc Test*  Vehicle vs Go6983  P=0.577  Vehicle vs Cmp101  P=0.239  Vehicle vs Go6983+Cmp101  P=0.923  Go6983 vs Cmp101  P=0.901  Go6983 vs Go6983+Cmp101  0.267  Cmp101 vs Go6983+Cmp101  P=0.088 | N/A  N/A  N/A  N/A | N/A |
| 5f | L149P-hβarr1: hGhrelin C/R  (Δ net BRET; % WT-vehicle) | vehicle  Go6983  Cmp101  Go6983+Cmp101 | N/A  N/A  N/A  N/A | N/A | N/A | 7.902±0.096  7.742±0.264  7.655±0.396  6.846±0.620 | F(3,336)=0.485  P=0.692 |
| 5g | L149P-hβarr1: hGhrelin E_max_  (E_max_; % WT-vehicle) | vehicle  Go6983  Cmp101  Go6983+Cmp101 | 44.760±1.784  39.951±4.264  15.288±2.964  8.834±3.284 | F(3,344)=35.05  P<0.0001**** | *Tukey’s Post Hoc Test*  Vehicle vs Go6983  P=0.605  Vehicle vs Cmp101  P<0.0001****  Vehicle vs Go6983+Cmp101  P<0.0001****  Go6983 vs Cmp101  P<0.0001****  Go6983 vs Go6983+Cmp101  P<0.0001****  Cmp101 vs Go6983+Cmp101  P=0.549 | N/A  N/A  N/A  N/A | N/A |
| 5i | WT-hβarr2: Constitutive  (Δ net BRET; % WT-vehicle) | vehicle  Go6983  Cmp101  Go6983+Cmp101 | 100±13.03  101.3±11.22  110.8±11.41  58.03±4.20 | F(3,68)=4.133  P=0.009** | *Tukey’s Post Hoc Test*  Vehicle vs Go6983  P=0.999  Vehicle vs Cmp101  P=0.901  Vehicle vs Go6983+Cmp101  P=0.044*  Go6983 vs Cmp101  P=0.931  Go6983 vs Go6983+Cmp101  P=0.035*  Cmp101 vs Go6983+Cmp101  P=0.011* | N/A  N/A  N/A  N/A | N/A |
| 5j | WT-hβarr2: hGhrelin C/R  (Δ net BRET; % WT-vehicle) | vehicle  Go6983  Cmp101  Go6983+Cmp101 | N/A  N/A  N/A  N/A | N/A | N/A | 7.791±0.071  7.758±0.079  7.904±0.098  UD | F(3,332)=0.294  P=0.829 |
| 5k | WT-hβarr2: hGhrelin E_max_  (E_max_; % WT-vehicle) | vehicle  Go6983  Cmp101  Go6983+Cmp101 | 100±3.555  74.813±3.072  32.099±1.951  18.204±1.603 | F(3,344)=144.7  P<0.0001**** | *Tukey’s Post Hoc Test*  Vehicle vs Go6983  P<0.0001****  Vehicle vs Cmp101  P<0.0001****  Vehicle vs Go6983+Cmp101  P<0.0001****  Go6983 vs Cmp101  P<0.0001****  Go6983 vs Go6983+Cmp101  P<0.0001****  Cmp101 vs Go6983+Cmp101  P=0.042* | N/A  N/A  N/A  N/A | N/A |
| 5l | L149P-hβarr2: Constitutive  (Δ net BRET; % WT-vehicle) | vehicle  Go6983  Cmp101  Go6983+Cmp101 | 36.88±5.949  42.97±4.405  56.35±5.796  28.49±4.397 | F(3,68)=4.525  P=0.005** | N/A | N/A  N/A  N/A  N/A | N/A |
| 5m | L149P-hβarr2: hGhrelin C/R  (Δ net BRET; % WT-vehicle) | vehicle  Go6983  Cmp101  Go6983+Cmp101 | N/A  N/A  N/A  N/A | N/A | N/A | 7.780±0.077  7.732±0.111  7.812±0.126  7.348±0.223 | F(3,336)=0.505  P=0.678 |
| 5n | L149P-hβarr2: hGhrelin E_max_  (E_max_; % WT-vehicle) | vehicle  Go6983  Cmp101  Go6983+Cmp101 | 72.894±2.365  67.237±3.149  31.022±1.693  18.463±1.968 | F(3,344)=107.8  P<0.0001**** | *Tukey’s Post Hoc Test*  Vehicle vs Go6983  P=0.326  Vehicle vs Cmp101  P<0.0001****  Vehicle vs Go6983+Cmp101  P<0.0001****  Go6983 vs Cmp101  P<0.0001****  Go6983 vs Go6983+Cmp101  P<0.0001****  Cmp101 vs Go6983+Cmp101  P=0.014* | N/A  N/A  N/A  N/A | N/A |
| 5p | WT-hβarr1: Constitutive  (Δ net BRET; % WT control) | control  GRK2^K220R^  GRK2i | 100±4.243  92.86±4.884  111.8±4.820 | F(2,25)=3.592  P=0.042* | *Tukey’s Post Hoc Test*  control vs GRK2^K220R^  P=0.588  Control vs GRK2i  P=0.157  GRK2^K220R^ vs GRK2i  P=0.044* | N/A  N/A  N/A  N/A | N/A |
| 5q | WT-hβarr1: hGhrelin C/R  (Δ net BRET; % WT control) | control  GRK2^K220R^  GRK2i | N/A  N/A  N/A  N/A | N/A | N/A | 7.755±0.076  8.065±0.371  7.661±0.137 | F(2,159)=0.515  P=0.598 |
| 5r | WT-hβarr1: hGhrelin E_max_  (E_max_; % WT control) | control  GRK2^K220R^  GRK2i | 100±3.164  18.191±3.005  81.938±4.959 | F(2,165)=90.39  P<0.0001**** | *Tukey’s Post Hoc Test*  control vs GRK2^K220R^  P<0.0001****  Control vs GRK2i  P=0.002**  GRK2^K220R^ vs GRK2i  P<0.0001**** | N/A  N/A  N/A  N/A | N/A |
| 5s | L149P-hβarr1: Constitutive  (Δ net BRET; % WT control) | control  GRK2^K220R^  GRK2i | 96.84±8.101  123.6±5.143  123.6±5.241 | F(2,25)=5.155  P=0.013* | *Tukey’s Post Hoc Test*  control vs GRK2^K220R^  P=0.054  Control vs GRK2i  P=0.022*  GRK2^K220R^ vs GRK2i  P=0.999 | N/A  N/A  N/A  N/A | N/A |
| 5t | L149P-hβarr1: hGhrelin C/R  (Δ net BRET; % WT control) | control  GRK2^K220R^  GRK2i | N/A  N/A  N/A  N/A | N/A | N/A | 7.705±0.128  8.072±0.591  7.512±0.135 | F(2,159)=0.759  P=0.469 |
| 5u | L149P-hβarr1: hGhrelin E_max_  (E_max_; % WT control) | control  GRK2^K220R^  GRK2i | 51.23±2.733  17.866±4.097  48.672±2.884 | F(2,165)=27.46  P<0.0001**** | *Tukey’s Post Hoc Test*  control vs GRK2^K220R^  P<0.0001****  Control vs GRK2i  P=0.802  GRK2^K220R^ vs GRK2i  P<0.0001**** | N/A  N/A  N/A  N/A | N/A |
| 5w | WT-hβarr2: Constitutive  (Δ net BRET; % WT control) | control  GRK2^K220R^  GRK2i | 100±5.858  -0.839±13.77  80.61±6.726 | F(2,31)=37.05  P<0.0001**** | *Tukey’s Post Hoc Test*  control vs GRK2^K220R^  P<0.0001****  Control vs GRK2i  P=0.230  GRK2^K220R^ vs GRK2i  P<0.0001**** | N/A  N/A  N/A  N/A | N/A |
| 5x | WT-hβarr2: hGhrelin C/R  (Δ net BRET; % WT control) | control  GRK2^K220R^  GRK2i | N/A  N/A  N/A  N/A | N/A | N/A | 7.677±0.062  7.996±0.095  7.643±0.176 | F(2,192)=2.806  P=0.063 |
| 5y | WT-hβarr2: hGhrelin E_max_  (E_max_; % WT control) | control  GRK2^K220R^  GRK2i | 100±2.820  43.184±3.130  73.295±2.983 | F(2,195)=85.65  P<0.0001**** | *Tukey’s Post Hoc Test*  control vs GRK2^K220R^  P<0.0001****  Control vs GRK2i  P<0.0001****  GRK2^K220R^ vs GRK2i  P<0.0001**** | N/A  N/A  N/A  N/A | N/A |
| 5z | L149P-hβarr2: Constitutive  (Δ net BRET; % WT control) | control  GRK2^K220R^  GRK2i | 77.91±7.094  53.72±8.731  85.25±6.666 | F(2,31)=4.015  P=0.028 | *Tukey’s Post Hoc Test*  control vs GRK2^K220R^  P=0.077  Control vs GRK2i  P=0.761  GRK2^K220R^ vs GRK2i  P=0.029* | N/A  N/A  N/A  N/A | N/A |
| 5aa | L149P-hβarr2: hGhrelin C/R  (Δ net BRET; % WT control) | control  GRK2^K220R^  GRK2i | N/A  N/A  N/A  N/A | N/A | N/A | 7.759±0.083  7.923±0.118  7.599±0.086 | F(2,192)=0.800  P=0.450 |
| 5bb | L149P-hβarr2: hGhrelin E_max_  (E_max_; % WT control) | control  GRK2^K220R^  GRK2i | 69.715±2.298  42.071±2.758  66.708±1.646 | F(2,195)=31.07  P<0.0001**** | *Tukey’s Post Hoc Test*  control vs GRK2^K220R^  P<0.0001****  Control vs GRK2i  P=0.649  GRK2^K220R^ vs GRK2i  P<0.0001**** | N/A  N/A  N/A  N/A | N/A |
| 5dd | GRK2^WT^: Constitutive  (Δ net BRET; % WT-vehicle) | WT-vehicle  WT-GRK2i  L149P-vehicle  L149P-GRK2i | 100±3.769  90.079±4.723  100.884±5.320  104.322±7.384 | *Interaction:*  F(1,44)=1.496  P=0.227  *GRK2i:*  F(1,44)=0.352  P=0.555  *Receptor:*  F(1,44)=1.918  P=0.173 | N/A | N/A  N/A  N/A  N/A | N/A |
| 5ee | GRK2^WT^: hGhrelin C/R  (Δ net BRET; % WT-vehicle) | WT-vehicle  WT-GRK2i  L149P-vehicle  L149P-GRK2i | N/A  N/A  N/A  N/A | N/A | N/A | 7.546±0.040  7.546±0.059  7.652±0.059  7.610±0.069 | F(3,272)=0.532  P=0.660 |
| 5ff | GRK2^WT^: hGhrelin E_max_  (E_max_; % WT-vehicle) | WT-vehicle  WT-GRK2i  L149P-vehicle  L149P-GRK2i | 100±1.876  90.567±2.354  86.828±2.540  86.530±2.806 | *Interaction:*  F(1,284)=3.592  P=0.059  *GRK2i:*  F(1,284)=4.074  P=0.044*  *Receptor:*  F(1,284)=12.71  P=0.0004*** | *Tukey’s Post Hoc Test*  WT-vehicle vs L149P-vehicle  P=0.0008***  WT-vehicle vs WT-GRK2i  P=0.030*  WT-vehicle vs L149P-GRK2i  P=0.0006***  L149P-vehicle vs WT-GRK2i  P=0.694  L149P-vehicle vs L149P-GRK2i  P=0.999  WT-GRK2i vs L149P-GRK2i  P=0.639 | N/A  N/A  N/A  N/A | N/A |
| 5gg | GRK2^ΔV661-ct^: Constitutive  (Δ net BRET; % WT) | WT  L149P | 100±4.048  98.76±4.767 | T(22)=0.197  P=0.842 | N/A | N/A  N/A | N/A |
| 5hh | GRK2^ΔV661-ct^: hGhrelin C/R  (Δ net BRET; % WT) | WT  L149P | N/A  N/A  N/A  N/A | N/A | N/A | 7.467±0.317  7.429±0.284 | UD* |
| 5ii | GRK2^ΔV661-ct^: hGhrelin E_max_  (E_max_; % WT) | WT  L149P | 100±10.076  117.03±12.935 | T(142)=1.039  P=0.300 | N/A | N/A  N/A  N/A  N/A | N/A |

N/A: Not Applicable; UD: Undeterminable; UD*: Undeterminable due to different curve fits (3- vs 4-parameters).

**Supplementary Table 5.** Data and statistics from main **Figure 4**.

| ***Figure/Panel*** | ***Description & Units*** | ***Condition/***  ***Treatment*** | ***Mean ± SEM*** | ***t/F-Statistic &***  ***P-value*** | ***Multiple Comparisons*** | ***pEC_50_ ± SEM*** | ***F-Statistic &***  ***P-value*** |
| --- | --- | --- | --- | --- | --- | --- | --- |
| 5b | WT-hβarr1: Constitutive  (Δ net BRET; % parental-WT) | Parental  ΔGRK2/3/5/6  ΔGRK2/3  ΔGRK5/6 | 100±7.483  97.20±8.395  109.2±10.90  93.71±5.7 | F(3,44)=0.630  P=0.599 | N/A | N/A  N/A  N/A  N/A | N/A |
| 5c | WT-hβarr1: hGhrelin C/R  (Δ net BRET; % parental-WT) | Parental  ΔGRK2/3/5/6  ΔGRK2/3  ΔGRK5/6 | N/A  N/A  N/A  N/A | N/A | N/A | 7.703±0.089  UD  UD  7.777±0.143 | F(3,270)=0.122  P=0.946 |
| 5d | WT-hβarr1: hGhrelin E_max_  (Δ net BRET; % parental-WT) | Parental  ΔGRK2/3/5/6  ΔGRK2/3  ΔGRK5/6 | 100±3.704  5.076±2.543  14.033±4.691  99.665±5.906 | F(3,278)=136.9  P<0.0001**** | *Tukey’s Post Hoc Test*  Parental vs ΔGRK2/3/5/6  P<0.0001****  Parental vs ΔGRK2/3  P<0.0001****  Parental vs ΔGRK5/6  P>0.999  ΔGRK2/3/5/6 vs ΔGRK2/3  P=0.496  ΔGRK2/3/5/6 vs ΔGRK5/6  P<0.0001****  ΔGRK2/3 vs ΔGRK5/6  P<0.0001**** | N/A  N/A  N/A  N/A | N/A |
| 5e | L149P-hβarr1: Constitutive  (Δ net BRET; % parental-WT) | Parental  ΔGRK2/3/5/6  ΔGRK2/3  ΔGRK5/6 | 103.4±5.975  95.41±5.632  104.5±10.13  112.1±5.195 | F(3,44)=0.950  P=0.424 | N/A | N/A  N/A  N/A  N/A | N/A |
| 5f | L149P-hβarr1: hGhrelin C/R  (Δ net BRET; % parental-WT) | Parental  ΔGRK2/3/5/6  ΔGRK2/3  ΔGRK5/6 | N/A  N/A  N/A  N/A | N/A | N/A | 7.419±0.431  UD  UD  7.259±0.620 | F(3,276)=0.842  P=0.471 |
| 5g | L149P-hβarr1: hGhrelin E_max_  (Δ net BRET; % parental-WT) | Parental  ΔGRK2/3/5/6  ΔGRK2/3  ΔGRK5/6 | 31.271±4.850  -2.820±2.243  0.000±0.000^#^  19.797±6.059 | F(3,284)=16.26  P<0.0001**** | *Tukey’s Post Hoc Test*  Parental vs ΔGRK2/3/5/6  P<0.0001****  Parental vs ΔGRK2/3  P<0.0001****  Parental vs ΔGRK5/6  P=0.187  ΔGRK2/3/5/6 vs ΔGRK2/3  P=0.960  ΔGRK2/3/5/6 vs ΔGRK5/6  0.0005***  ΔGRK2/3 vs ΔGRK5/6  P=0.003** | N/A  N/A  N/A  N/A | N/A |
| 5i | WT-hβarr2: Constitutive  (Δ net BRET; % parental-WT) | Parental  ΔGRK2/3/5/6  ΔGRK2/3  ΔGRK5/6 | 100±4.707  97.54±3.483  102.1±5.582  88.19±3.219 | F(3,44)=1.989  P=0.129 | N/A | N/A  N/A  N/A  N/A | N/A |
| 5j | WT-hβarr2: hGhrelin C/R  (Δ net BRET; % parental-WT) | Parental  ΔGRK2/3/5/6  ΔGRK2/3  ΔGRK5/6 | N/A  N/A  N/A  N/A | N/A | N/A | 7.962±0.064  UD  7.475±0.573  7.900±0.076 | F(3,276)=0.537  P=0.656 |
| 5k | WT-hβarr2: hGhrelin E_max_  (Δ net BRET; % parental-WT) | Parental  ΔGRK2/3/5/6  ΔGRK2/3  ΔGRK5/6 | 100±1.966  -1.031±1.773  14.023±2.813  124.150±3.859 | F(3,284)=470.9  P<0.0001**** | *Tukey’s Post Hoc Test*  Parental vs ΔGRK2/3/5/6  P<0.0001****  Parental vs ΔGRK2/3  P<0.0001****  Parental vs ΔGRK5/6  P<0.0001****  ΔGRK2/3/5/6 vs ΔGRK2/3  0.001**  ΔGRK2/3/5/6 vs ΔGRK5/6  P<0.0001****  ΔGRK2/3 vs ΔGRK5/6  P<0.0001**** | N/A  N/A  N/A  N/A | N/A |
| 5l | L149P-hβarr2: Constitutive  (Δ net BRET; % parental-WT) | Parental  ΔGRK2/3/5/6  ΔGRK2/3  ΔGRK5/6 | 88.89±4.654  95.50±8.691  100.1±11.10  75.32±4.254 | F(3,44)=1.957  P=0.134 | N/A | N/A  N/A  N/A  N/A | N/A |
| 5m | L149P-hβarr2: hGhrelin C/R  (Δ net BRET; % parental-WT) | Parental  ΔGRK2/3/5/6  ΔGRK2/3  ΔGRK5/6 | N/A  N/A  N/A  N/A | N/A | N/A | 7.971±0.148  7.660±0.946  7.926±0.536  7.531±0.173 | F(3,264)=1.094  P=0.352 |
| 5n | L149P-hβarr2: hGhrelin E_max_  (Δ net BRET; % parental-WT) | Parental  ΔGRK2/3/5/6  ΔGRK2/3  ΔGRK5/6 | 45.841±2.490  11.146±2.869  16.598±2.405  30.166±2.197 | F(3,272)=38.69  P<0.0001**** | *Tukey’s Post Hoc Test*  Parental vs ΔGRK2/3/5/6  P<0.0001****  Parental vs ΔGRK2/3  P<0.0001****  Parental vs ΔGRK5/6  P<0.0001****  ΔGRK2/3/5/6 vs ΔGRK2/3  P=0.429  ΔGRK2/3/5/6 vs ΔGRK5/6  P<0.0001****  ΔGRK2/3 vs ΔGRK5/6  P=0.0006*** | N/A  N/A  N/A  N/A | N/A |
| 5p | GRK5^YFP^:  Constitutive Activity  (Δ net BRET) | WT  L149P | 100±20.72  103.8±22.36 | *Normality Tests*  *Shapiro-Wilk*  P<0.0001****  Kolmogorov-Smirnov  P<0.0001****  *Non-Parametric*  *Mann-Whitney* U = 68  P=0.842 | N/A | N/A  N/A  N/A  N/A | N/A |
| 5q | GRK5^YFP^:  hGhrelin C/R^#^  (net BRET) | WT-basal  WT-hGhr^10pM^  WT-hGhr^100pM^  WT-hGhr^1nM^  WT-hGhr^10nM^  WT-hGhr^100nM^  WT-hGhr^1μM^  L149P-basal  L149P-hGhr^10pM^  L149P-hGhr^100pM^  L149P-hGhr^1nM^  L149P-hGhr^10nM^  L149P-hGhr^100nM^  L149P-hGhr^1μM^ | 1.206±0.023  1.212±0.020  1.205±0.018  1.209±0.016  1.209±0.018  1.220±0.018  1.224±0.019  1.236±0.014  1.235±0.010  1.236±0.010  1.224±0.016  1.222±0.013  1.259±0.014  1.254±0.015 | *Interaction:*  F(6,132)=1.00  P=0.425  *hGhrelin:*  F(3.5,79.0)=4.89  P=0.002**  *Receptor:*  F(1,22)=1.42  P=0.245 | N/A | N/A  N/A  N/A  N/A | N/A |
| 5r | GRK6^YFP^:  Constitutive Activity  (Δ net BRET) | WT  L149P | 100±14.39  163.3±27.34 | *Normality Tests*  *Shapiro-Wilk*  P<0.0004***  Kolmogorov-Smirnov  P<0.0001****  *Non-Parametric*  Mann-Whitney U = 27  P=0.008** | N/A | N/A  N/A  N/A  N/A | N/A |
| 5s | GRK6^YFP^:  hGhrelin C/R^#^  (net BRET) | WT-basal  WT-hGhr^10pM^  WT-hGhr^100pM^  WT-hGhr^1nM^  WT-hGhr^10nM^  WT-hGhr^100nM^  WT-hGhr^1μM^  L149P-basal  L149P-hGhr^10pM^  L149P-hGhr^100pM^  L149P-hGhr^1nM^  L149P-hGhr^10nM^  L149P-hGhr^100nM^  L149P-hGhr^1μM^ | 1.466±0.011  1.456±0.011  1.459±0.007  1.466±0.007  1.449±0.011  1.462±0.013  1.465±0.010  1.586±0.008  1.581±0.008  1.589±0.010  1.571±0.012  1.572±0.013  1.575±0.010  1.604±0.012 | *Interaction:*  F(6,132)=1.811  P=0.101  *hGhrelin:*  F(4.7,103.9)=3.2  P=0.010*  *Receptor:*  F(1,22)=95.67  P<0.0001**** | *Tukey’s Post Hoc Test*  Basal (WT vs L149)  P<0.0001****  hGhr^10pM^ (WT vs L149)  P<0.0001****  hGhr^100pM^ (WT vs L149)  P<0.0001****  hGhr^1nM^ (WT vs L149)  P<0.0001****  hGhr^10nM^ (WT vs L149)  P<0.0001****  hGhr^100nM^ (WT vs L149)  P<0.0001****  hGhr^1μM^ (WT vs L149)  P<0.0001**** | N/A  N/A  N/A  N/A | N/A |

N/A: Not Applicable; UD: Undeterminable; ^#^ Unable to fit curve

**Supplementary Table 6.** Data and statistics from main **Figure 5**.

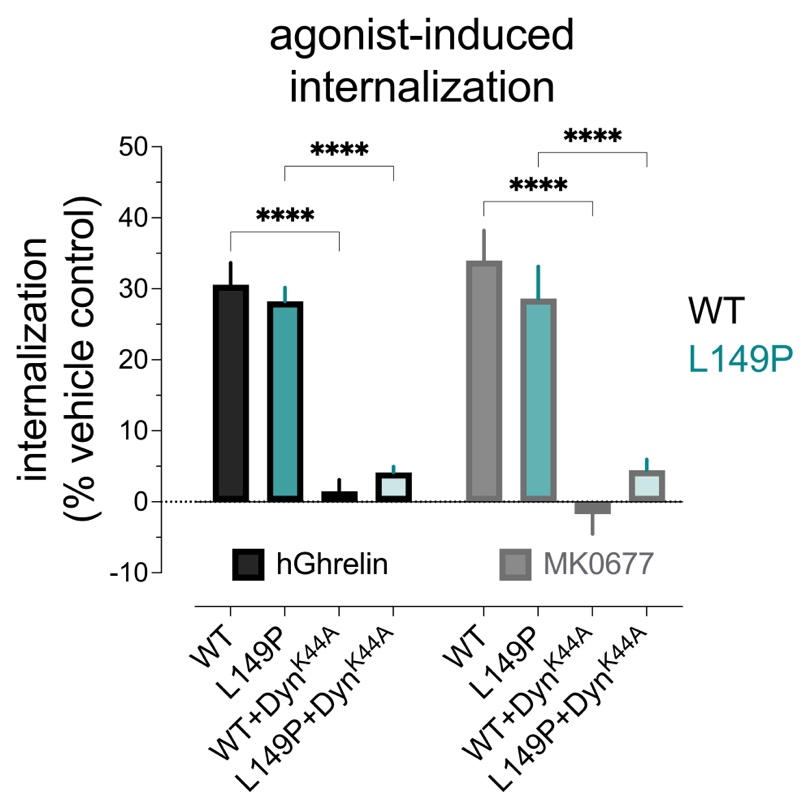
**SUPPLEMENTARY FIGURES & LEGENDS**

**Supplementary Figure 1. Agonist-stimulated, dynamin-dependent internalization of the WT and L149P receptors.** Percent agonist-induced internalization of the WT (black) or L149P (teal) receptors in response to hGhrelin (black border) or MK0677 (grey border) derived from *Figure 1b*. To quantify internalization (loss of surface expression), the data for each condition were normalized to their respective vehicle-treated controls. All data represent the pooled mean ± SEM from 3 independent experiments run in quadruplicate or sextuplicate.

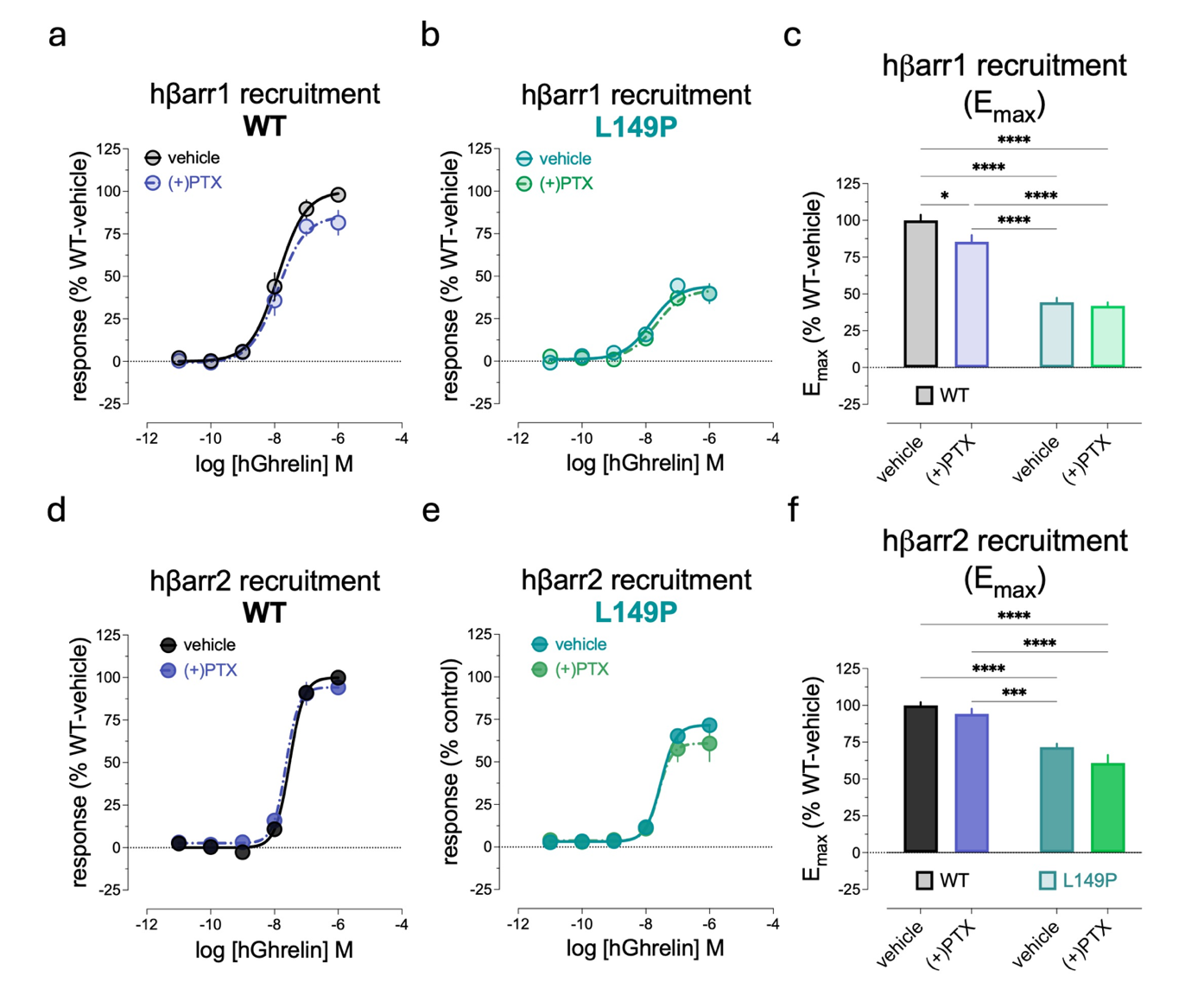

**Supplementary Figure 2. Agonist-stimulated hβarr1/2 recruitment to L149P is Gα_i/o_-independent.** **(a)** hGhrelin (C/R)-induced hβarr1 recruitment to WT following vehicle (grey) or overnight PTX (200ng/mL; light blue) pretreatment. **(b)** hGhrelin (C/R)-induced hβarr1 recruitment to L149P following vehicle (light teal) or PTX pretreatment. **(c)** hGhrelin E_max_ (derived from *Panels a-b*) of hβarr1 recruitment to WT (*left*) and L149P (*right*) ± PTX. **(d)** hGhrelin (C/R)-induced hβarr2 recruitment to WT following vehicle (black) or overnight PTX (200ng/mL; dark blue) pretreatment. **(e)** hGhrelin (C/R)-induced hβarr2 recruitment to L149P following vehicle (dark teal) or PTX pretreatment. **(f)** hGhrelin E_max_ (derived from *Panels d-e*) of hβarr2 recruitment to WT (*left*) and L149P (*right*) ± PTX. All data represent the pooled mean ± SEM from 3 independent experiments run in duplicate.
